# *In vivo* Interactions between Myosin XI, Vesicles, and Filamentous Actin Are Fast and Transient

**DOI:** 10.1101/624361

**Authors:** Jeffrey P. Bibeau, Fabienne Furt, S. Iman Mousavi, James L. Kingsley, Max F. Levine, Erkan Tüzel, Luis Vidali

## Abstract

The apical actin cytoskeleton and active membrane trafficking machinery are essential in driving polarized cell growth. To better understand the interactions between myosin XI, vesicles, and actin filament *in vivo*, we performed Fluorescence Recovery After Photobleaching (FRAP) and showed that the dynamics of myosin XIa at the tip are actin-dependent and that approximately half of myosin XI is bound to vesicles in the cell. To obtain single particle information, we used Variable Angle Epifluorescence Microscopy (VAEM) in *Physcomitrella patens* protoplasts to demonstrate that myosin XIa and VAMP72-labeled vesicles localize in time and space for periods lasting only a few seconds. Using tracking data with Hidden Markov Modeling (HMM), we showed that myosin XIa and VAMP72-labeled vesicles exhibit short runs of actin-dependent directed transport. We also found that the interaction of myosin XI with vesicles is short lived. Together, this bound fraction, fast off-rate, and short run lengths are expected to be critical for the dynamic oscillations observed at the cell apex, and may be vital for the regulation and recycling of the exocytosis machinery; while simultaneously promoting the vesicle focusing and secretion at the tip, necessary for cell wall expansion.

## Introduction

Polarized cell growth, a mechanism by which plasma membrane and cell wall material are deposited to a defined region of the cell, is widespread across the plant kingdom. As a result, plant cells exhibit a large variety of different shapes well designed to achieve various functions (Geitmann and Ortega, 2009; Szymanski and Cosgrove, 2009). Tip growth is an extreme form of polarized cell growth where the cell expands only in one direction generating elongated cells that can be hundreds of micrometers long (Hepler et al., 2001; Menand et al., 2007). This specialized form of growth is particularly well suited for pollen tubes to reach the ovule during sexual reproduction (Cole and Fowler, 2006; Hepler et al., 2001), for root hairs to enhance uptake nutrients and water from the soil (Gilroy and Jones, 2000; Hepler et al., 2001), and for bryophytes for land colonization (Heckman et al., 2001; Kenrick and Crane, 1997). This process has been extensively studied in pollen tubes from lily, tobacco, and *Arabidopsis thaliana* and in root hairs from *A. thaliana* and protonemal cells from *Physcomitrella patens*. It is well known that tip growth requires a dynamic actin cytoskeleton network as well as active membrane trafficking machinery (Cole and Fowler, 2006; Gilroy and Jones, 2000; Heckman et al., 2001; Hepler et al., 2001; Kenrick and Crane, 1997). Modeling evidence suggests that endomembrane vesicles must be focused at the tip of the cell by F-actin to overcome diffusion limitations and sustain growth (Bibeau et al., 2018). Because the wall material required for growth far exceeds the necessary membrane material, endocytosis occurs to recycle the excess plasma membrane. However, how these different types of machineries are coordinated with the actin cytoskeleton is still unknown. To better understand how tip growing cells self-organize their cytoplasmic components to achieve and maintain polarized growth, it is critical to determine the function of each component separately as well as their interactions. Because of this limited information, quantitative modeling efforts must coarse grain the function of cytoskeleton as directed secretion fluxes (Campàs and Mahadevan, 2009; Dumais et al., 2006; Fayant et al., 2010; Kroeger et al., 2011; Luo et al., 2017; Rojas et al., 2011). Although these fluxes are a good first approximation, they provide little insight into the mechanisms by which the cytoskeleton functions. Modeling efforts that incorporate the true interactions of the cytoskeleton, its motors, and cargo will help answer questions regarding the establishment and maintenance of polarized secretion.

Several studies have shown that the actin-based molecular motor myosin XI is required for tip growth in plants. Silencing of the two myosin XI genes—Myosin XIa and b—in the moss *Physcomitrella patens* resulted in round cells where tip growth was completely abolished (Vidali et al., 2010). Dominant negative inhibition and gene knockout approaches suggested that among the 13 isoforms of myosin XIa present in *A. thaliana*, myosin XI-K is a primary contributor for root hair elongation by tip growth (Ojangu et al., 2007; Park and Nebenfuhr, 2013; Peremyslov et al., 2008; Prokhnevsky et al., 2008); while myosin XIc1 and c2 are important for pollen tube growth (Madison et al., 2015). In the last decade, it has been well established that myosin XIs are responsible for the motility of large organelles in higher plants, even though their relative contribution and interdependency relationships are not fully understood (Avisar et al., 2012; Avisar et al., 2009; Avisar et al., 2008; Griffing et al., 2014; Henn and Sadot, 2014; Madison and Nebenfuhr, 2013; Peremyslov et al., 2008; Peremyslov et al., 2010, 2010, 2010, 2010, 2010; Prokhnevsky et al., 2008; Sparkes et al., 2008; Ueda et al., 2010; Vick and Nebenfuhr, 2012). However, the organelle motility was not impaired in the *myosin xi-k* knockout mutants which display anormal root hair elongation (Prokhnevsky et al., 2008). In addition, the apical localization of myosin XIa in moss caulonemal cells does not correlate with the localization observed for the large organelles in other plants (Furt et al., 2012; Vidali et al., 2010). Therefore, the function fulfilled by myosin XI that is essential to achieve and/or maintain tip growth in plants is still unknown.

Numerous studies have reported that myosin Vs, which are the homologues of myosin XIs in animals and yeast, are involved not only in large organelle motility but also in secretory, endocytic and recycling pathways via their ability to transport endomembrane vesicles (Fan et al., 2004; Hammer and Sellers, 2012; Lapierre et al., 2001; Li and Nebenfuhr, 2008; Lise et al., 2006; Nedvetsky et al., 2007; Ohyama et al., 2001; Pruyne et al., 1998; Rodriguez and Cheney, 2002; Roland et al., 2007; Schott et al., 1999; Volpicelli et al., 2002; Yan et al., 2005). Recent evidence suggests that myosin XIs could be involved in endomembrane vesicle transport in plants. Results from co-localization with vesicle markers (RabA4b and SCAMP2), fluorescence recovery after photobleaching, and biochemical co-fractionation with vesicle (RabA4b) and exocytic (Sec6) markers inferred that myosin XI-K is associated with endomembrane vesicles in both leaf cells and root hairs in *A. thaliana* (Park and Nebenfuhr, 2013; Peremyslov et al., 2012; Rybak et al., 2014). Using confocal microscopy combined with fluctuation cross-correlation analyses, we previously demonstrated that myosin XIa and VAMP72, an endomembrane vesicle marker, co-localize at the cell apex and are synchronized during tip growth (Furt et al., 2013). Surprisingly, we also showed that apical myosin XIa precedes F-actin during polarized growth of *P. patens* caulonemal cells (Furt et al., 2013). Pharmacological approaches using latrunculin B to depolymerize the actin filaments, further showed that myosin XIa stays associated with the VAMP72 marker at the apex of moss cells after disruption of the actin network (Furt et al., 2013). In addition, we observed the emergence of ectopic clusters of myosin XIa associated with VAMP72-labeled vesicles, which were then propelled through the cell via an actin-dependent manner when the actin network self-reorganizes (Furt et al., 2013).

Together, these results suggest the existence of a mechanism where myosin XI-associated endomembrane structures could organize the actin polymerization machinery. However, the apex of plant tip growing cells being densely occupied by vesicles (Bove et al., 2008; de Graaf et al., 2005; Kroeger et al., 2009; Ovecka et al., 2005; Parton et al., 2001; Preuss et al., 2004; Szumlanski and Nielsen, 2009; Zonia and Munnik, 2008), and the resolution of confocal laser scanning microscopes being diffraction limited, make it challenging to visualize single vesicles (~80 nm in diameter) in plant tip growing cells to uncover this mechanism. Therefore, the ability of myosin XIa to transport endomembrane vesicles and coordinate this transport with the actin polymerization machinery remains to be demonstrated, and whether this is the essential function fulfilled by myosin XIa to achieve polarized growth remains to be proved. Furthermore, there are no quantitative *in vivo* estimates of these binding interactions in plants. Specifically, the average on and off-rates of myosin XI and its cargo are not known. In addition, myosin XI run lengths and off-rates on actin filaments remain unknown. For these reasons, any quantitative insight into these interactions would shape the way the active transport machinery is viewed in plants. Furthermore, it would enable future modeling by constraining the large parameter space of the cytoskeleton in polarized growth.

In this study, we used fluorescence recovery after photobleaching (FRAP) (Loren et al., 2015; McNally, 2008) to demonstrate that myosin XIa recovers at the cell apex in an actin-dependent manner. With the actin depolymerizing agent Latrunculin B, we performed FRAP in the absence of actin filament and determined a range of effective dissociation constants for myosin XIa and VAMP72-labeled vesicles. To improve the precision of these measurements we used variable angle epifluorescence microscopy (VAEM). We showed that both myosin XIa and VAMP72 co-localize at the cortex of moss cells to highly dynamic punctate structures, whose motility depends on actin filaments. With particle tracking and Hidden Markov Modeling (HMM), we characterized the weak transient interactions between myosin XI, VAMP72-labeled vesicles, and filamentous actin. Our results are consistent with the hypothesis that myosin XIa coordinates the traffic machinery and the actin network dynamics to maintain polarized growth in moss cells.

## Results

### Apical Myosin XIa Recovers Faster than VAMP72-vesicels at the Cell Apex in an Actin-Dependent Manner

We previously showed that myosin XIa fluctuations anticipate F-actin’s at the apex of caulonemal cells while myosin XIa levels fluctuate in identical phase with VAMP72, an endomembrane vesicle marker (Furt et al., 2013). To explain this, we proposed a model where myosin XIa, via its association with endomembrane vesicles, coordinates the vesicular traffic and the F-actin polymerization-driven motility at the cell apex, to maintain polarized cell growth (Furt et al., 2013). This proposed model predicts that the dynamics of myosin XIs at the cell tip should exhibit some degree of coupling with filamentous actin. To verify this prediction, we conducted Fluorescence Recovery After Photobleaching (FRAP) experiments (Loren et al., 2015; McNally, 2008) of myosin XIa at the growing cell tip and shank (Figure 1A). Compared to the shank, myosin XIa at the tip exhibited an increased rate of fluorescence recovery. This suggests that F-actin, which concentrates at the cell tip, is coupled to an increased flow of myosin XIa. This result does not support a simple mechanism by which F-actin statically captures myosin XIa for increased accumulation at the cell tip, which should result in a slower myosin XIa recovery in FRAP.

**Figure 1.**
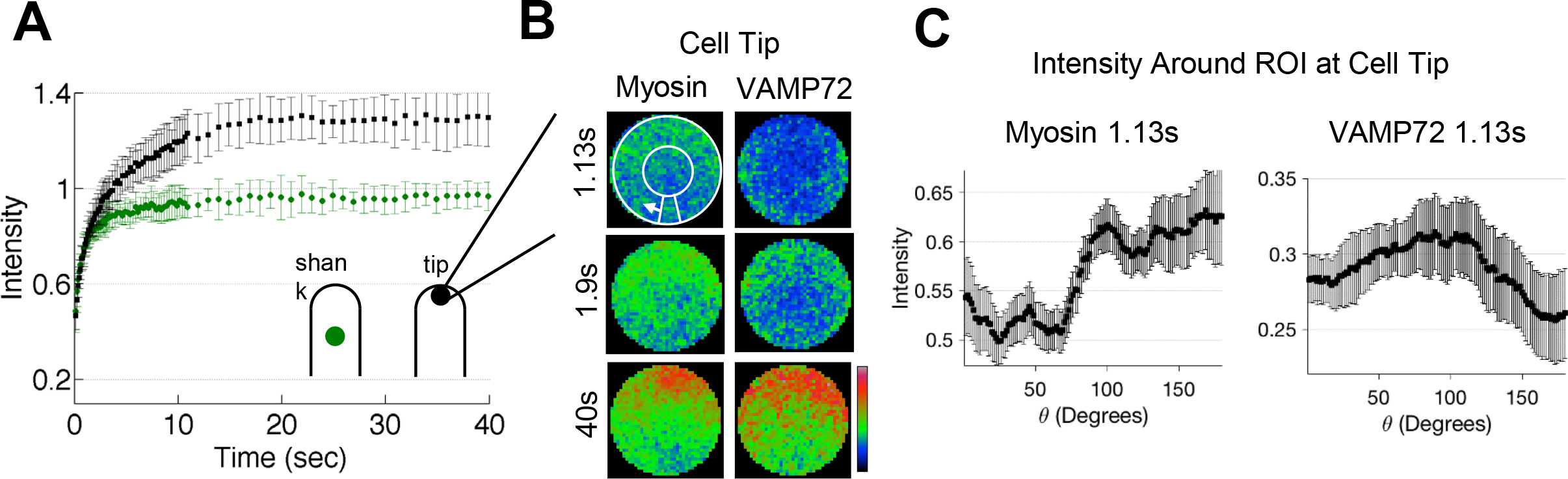
Apical Myosin XIa dynamics during polarized cell growth. **A)** Fluorescence recovery of 3mEGFP-myosin XIa at the cell shank (green circles) and the cell tip (black circles). n=7 and 10 for the shank and tip, respectively. (Error bars represent standard deviations). **B)** Cropped and frame averaged photobleaching ROI at the cell tip for cells expressing 3mEGFP-myosin XIa (left) and 3mCherry-VAMP72 (right). White circle and arrows indicate how the perimeter of the ROI was measured during the recovery. **C)** Intensity profile of 3mEGFP-myosin XIa (left) and 3mCherry-VAMP72 (right) along the perimeter of the ROI 1.13 s following photobleaching.

To better understand how F-actin coupling results in increased flow of myosin XIa at the cell tip, we analyzed its fluorescence recovery spatially (Figure 1B and Sup Movie 1). At early times during fluorescence recovery (a chosen time point of 1.13 s), we found that myosin XIa could be detected recovering at the cell apex before VAMP72-labeled vesicles (Figure 1B). Given that the bleach region is 4 μm in diameter, this fluorescence recovery would be consistent with myosin motor speeds of a few microns per second. To quantify the directionality of this recovery, we measured the fluorescence of myosin XIa along the perimeter of the photobleached region and found that myosin recovered first at the cell apex. (Figure 1C). This spatial recovery profile is not consistent with diffusion, which should recover from the shank towards the cell tip (Bibeau et al., 2018; Kingsley et al., 2018), or with VAMP72-vesicles, which recover along the actin cortex (Figure 1C). It is expected that the myosin XI recovery is a composite of vesicle transport, diffusion, and transport on actin independently of vesicles.

To determine if the observed myosin XIa fluorescence recovery was actin dependent, we treated growing cells with latrunculin B. Depolymerization of F-actin reduced the flow of myosin XIa at the cell tip (Figure 2A) and removed the myosin XIa dense apical spot (Figure 2B). Without F-actin, myosin XIa recovered from the cell shank toward the cell tip (Figure 2B, 2C and Sup Movie 2), in a pattern consistent with simple diffusion (Bibeau et al., 2018; Kingsley et al., 2018). This indicates that myosin XIa exhibits an actin-dependent apical recovery in the absence of latrunculin B treatment.

**Figure 2.**
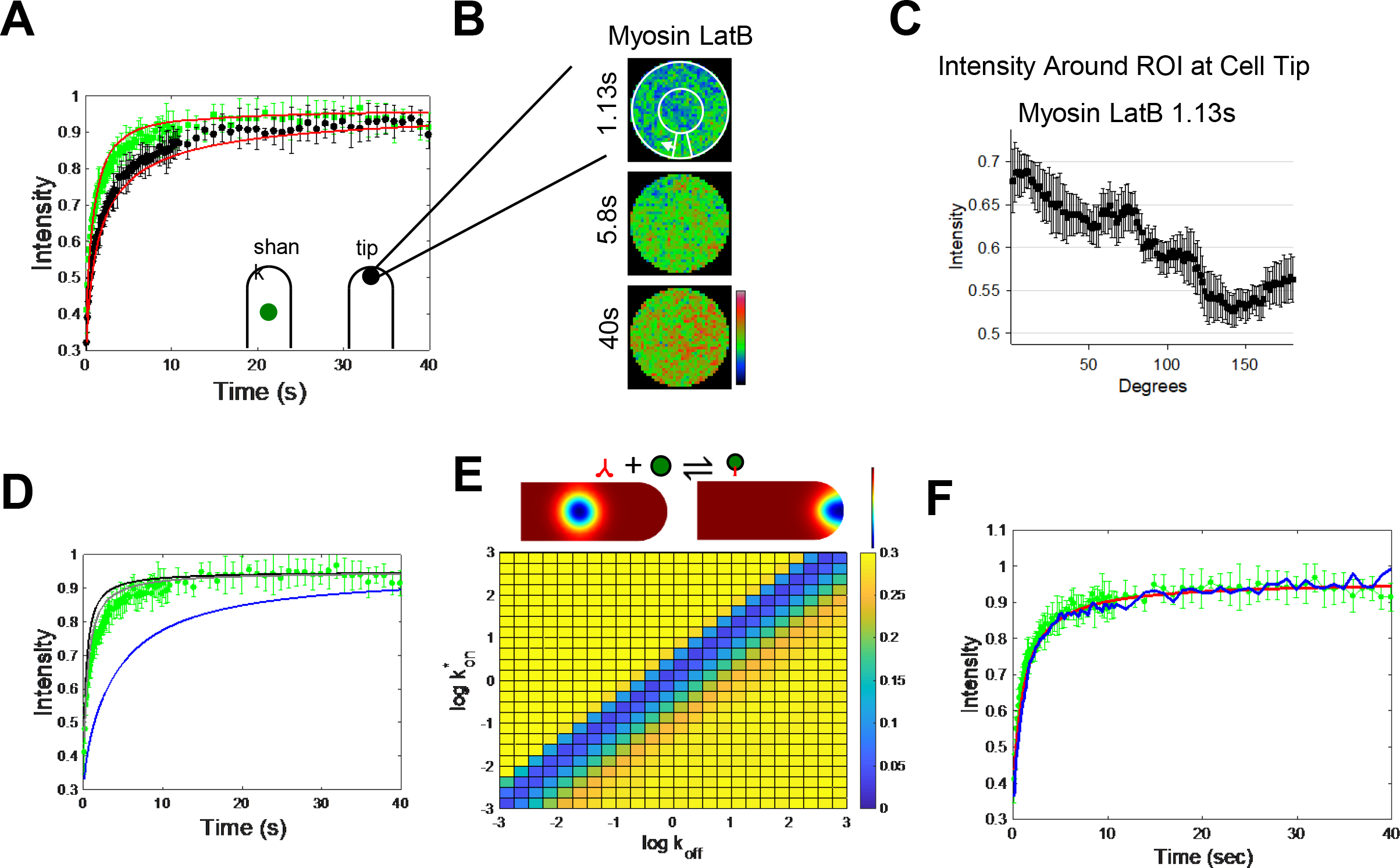
Latrunculin B treatment allows for characterization of myosin XIa and vesicle binding. **A)** Fluorescence recovery of 3mEGFP-myosin XIa in latrunculin B treated cells at the shank (green circles) and the tip (black circles). Red line indicates reaction diffusion FRAP model with best fit dissociation constant, K_d_. n=6 and 8 for the shank and tip, respectively. (error bars indicate standard deviations). **B)** Cropped and frame averaged photobleaching ROI at the cell tip for latrunculin B treated cells expressing 3mEGFP-myosin XIa. White circle and arrows indicate how the perimeter of the ROI was measured during the recovery. **C)** Intensity profile of 3mEGFP-myosin XIa along the perimeter of the ROI, 1.13 s following photobleaching. **D)** Fluorescence recovery of 3mEGFP-myosin XIa in latrunculin B treated cells at the shank (green circles) compared to theoretical myosin XIa diffusion where *D* = 2.3 (gray), 3.4 (black), or 0.29 μm^2^/s (blue). **E)** Example reaction-diffusion model of FRAP at the cell shank and tip 1.13 s following photobleaching (top). Red indicates concentrated pixels and blue indicates less concentrated pixels. *k_on_* and *k_off_* heat map of the sum of squared differences between the Comsol reaction-diffusion model at the cell shank and the experimental myosin XIa fluorescence recovery at the shank (bottom). Bright yellow indicates large differences and dark blue indicates small differences. **F)** Fluorescence recovery of 3mEGFP-myosin XIa in latrunculin B treated cells at the shank (green circles) and corresponding two-dimensional infinite-boundary reaction-diffusion model from (Kang et al., 2010) (red dotted lines). Blue line indicates the fluorescence recovery from Digital Confocal Microscopy Suite. Both simulations were run with 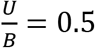.

### FRAP Reveals the Fraction of Myosin XI bound to VAMP72-vesicles

To determine the degree to which myosin XIa binds to VAMP72-labeled vesicles, we conducted FRAP experiments in the presence of latrunculin B (Figure 2A). By removing the actin cytoskeleton, we ensure that myosin XIa fluorescence recovery was a result of myosin XIa either bound or unbound to vesicles. Because myosin XI should diffuse more slowly when associated with vesicles, it is possible to infer the fraction of bound myosin XI by measuring the rate of the fluorescence recovery, if the diffusion coefficients are known. Since myosin XIa diffusion cannot be measured with FRAP, due to the potential of vesicle binding, we estimated a potential range of diffusion coefficients based on its similarity to the well-characterized myosin V (Krementsov et al., 2004; Li et al., 2004; Wang et al., 2004), and the measured viscosity of the moss cytoplasm (Kingsley et al., 2018) (see Methods). Since myosin V can be found in an open or closed configuration, we estimate that the myosin XI diffusion coefficient is between 2.3 and 3.4 μm^2^/s, respectively.

With both the myosin XI and vesicle (Bibeau et al., 2018) diffusion coefficients, we modeled the interaction as a reversible bimolecular reaction, *U* + *S* ⇌ *B*. Here *U* represents the concentration of unbound myosin XIa in the cytoplasm, *S* represents the concentration of available vesicle binding sites for myosin XIa, and *B* represents the concentration of myosin XIa bound to a vesicle receptor. We further assumed that the number of myosin XIa binding sites on a vesicle, *S(x,y,t),* is constant at equilibrium i.e *S(x,y,t)=S_eq_* (Kang and Kenworthy, 2008; Kang et al., 2010; Sprague and McNally, 2005). It then follows that we can write the following expression for the effective on-rate *k*_*on*_^*^=*S*_*eq*_*k*_*on*_ (Kang and Kenworthy, 2008; Kang et al., 2010; Sprague and McNally, 2005). Based on these assumptions, we can now model the process as a two-species reaction-diffusion system (Kang and Kenworthy, 2008; Kang et al., 2010) described by Eqs. 1 and 2, i.e.,

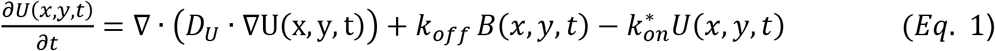

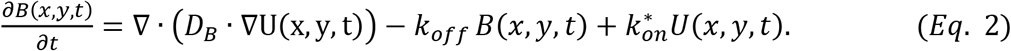

Here, *U(x,y,t)* and *B(x,y,t)* are the concentrations of unbound and bound myosin XIa, *D*_*U*_ and *D*_*B*_ are the estimated diffusion coefficients for unbound myosin XIa and VAMP72-labeled vesicles, ∇ is the gradient operator, and *k_off_* is the myosin XIa vesicle off-rate.

Although Eqs. 1 and 2 (Kang and Kenworthy, 2008; Kang et al., 2010; Sprague and McNally, 2005), can be solved analytically for simple boundary conditions, no analytical solution exists for our cell shape. Since cell shape has been shown to be important for modeling FRAP (Bibeau et al., 2018; Kingsley et al., 2018), we used the finite element modeling software Comsol (Comsol Inc, Stockholm, Sweden) to simulate FRAP in the presence of myosin vesicle binding (Figure 2E) (see Methods) and cell boundaries, to solve this system of equations. A parameter sweep across potential reaction on and off-rates (Figure 2E) demonstrates that we only have the experimental sensitivity to determine the effective dissociation constant, or more intuitively the ratio of bound to unbound myosin, *k*_*d*_^*^=*k*_*off*_/*k*_*on*_^*^=*U/B,* and not the individual rate constants. Using the Matlab (Mathworks, Natick, MA) liveLink Comsol Multiphysics module, we were able to iteratively find an effective dissociation constant that matched experimental myosin XIa fluorescence recoveries, *U/B= 0.8-0.5,* at both the tip and the shank (Figure 2A, see Methods for details). Here, the range in *U/B* reflects the possible open (0.8) and close (0.5) configurations of myosin XI. Through algebraic arrangement, *U/B* can be modified to be the fraction of myosin XI bound to a vesicle, which is approximately 55% (open) and 66%(close). Similar recovery curves were found by applying these best fit parameters to the two-dimensional infinite-boundary reaction-diffusion model from Kang *et al.* shank, (Figure 2F) (Kang et al., 2010) and Digital Confocal Microscopy Suite (DCMS) with reaction-diffusion (Kingsley et al., 2018)(Figure 2F).

### Myosin XIa and VAMP72 Puncta Exhibit Actin Dependent Dynamics at the Cortex of Moss Protonemal Cells

To further assess the dynamics of myosin XIa and vesicles on the particle level, we used Variable Angle Epifluorescence Microscopy (VAEM), which provides the resolution necessary to image single endomembrane vesicles (Konopka and Bednarek, 2008). We found, as expected that 3mEGFP-VAMP72 labeled highly dynamic small punctate structures at the cortex of apical caulonemal cells (Sup Figure S1A-top, Sup Movie 3). This is consistent with localization to endomembrane vesicles, and observations reported in protoplasts from *Arabidopsis* suspension cultured cells using the same probe (Uemura et al., 2004). We also showed that 3mEGFP-Myosin XIa localizes to similar dynamic punctate structures at the cortex of caulonemal cells (Sup Figure S1A-bottom – Sup Movie 3). Using the detection tool of the TrackMate plugin from ImageJ (see Experimental Procedures), we determined that the average density of myosin XIa-decorated structures is 0.49 +/− 0.04 per μm^2^, and is similar to that of VAMP72-labeled endomembrane vesicles with 0.52 +/− 0.01 per μm^2^ (Sup Figure S1B). These densities are slightly higher but in the same range as those reported for class II formin, a nucleator of actin filaments, in the same model system (van Gisbergen et al., 2012).

To evaluate if the motility of both myosin XIa- and VAMP72-labeled vesicles depends on actin, we imaged the apical caulonemal cells in presence or absence of 25 μM latrunculin B, which completely depolymerizes the actin filaments (Vidali et al., 2009). We tracked the movement of punctate structures labeled by myosin XIa and VAMP using TrackMate in presence or absence of the drug. At least three types of motion were observed: some punctate structures that move rapidly in and out of the imaging field, some move in a persistent linear manner, and some stay confined to an area (displacement below 0.5 μm). We only focused on punctate structures that display a directed movement for at least 12 consecutive frames (600 ms) and show a displacement higher than 1.2 μm. Examples of such persistent linear trajectories recorded for myosin XIa and VAMP72 are shown in figure 3A and B, respectively (Sup Movie 3). Over a period of 100 sec, we were able to detect 20.2 ± 3.1 and 13.3 ± 2.6 persistent linear trajectories per average imaged area per cell (40 μm^2^) for myosin XIa and VAMP72, respectively (Figures 3C and D). However, after treatment with latrunculin B, we found only 3.6 ± 0.2 and 0.8 ± 0.8 persistent linear trajectories per average imaged area per cell (40 μm^2^) for myosin XIa and VAMP72, respectively (Figures 3C and D). Such a large decrease in the number of persistent linear trajectories in the absence of actin filaments supports the hypothesis that myosin XIa- and VAMP72-labeled endomembrane vesicles move on actin filaments in apical caulonemal cells.

**Figure 3.**
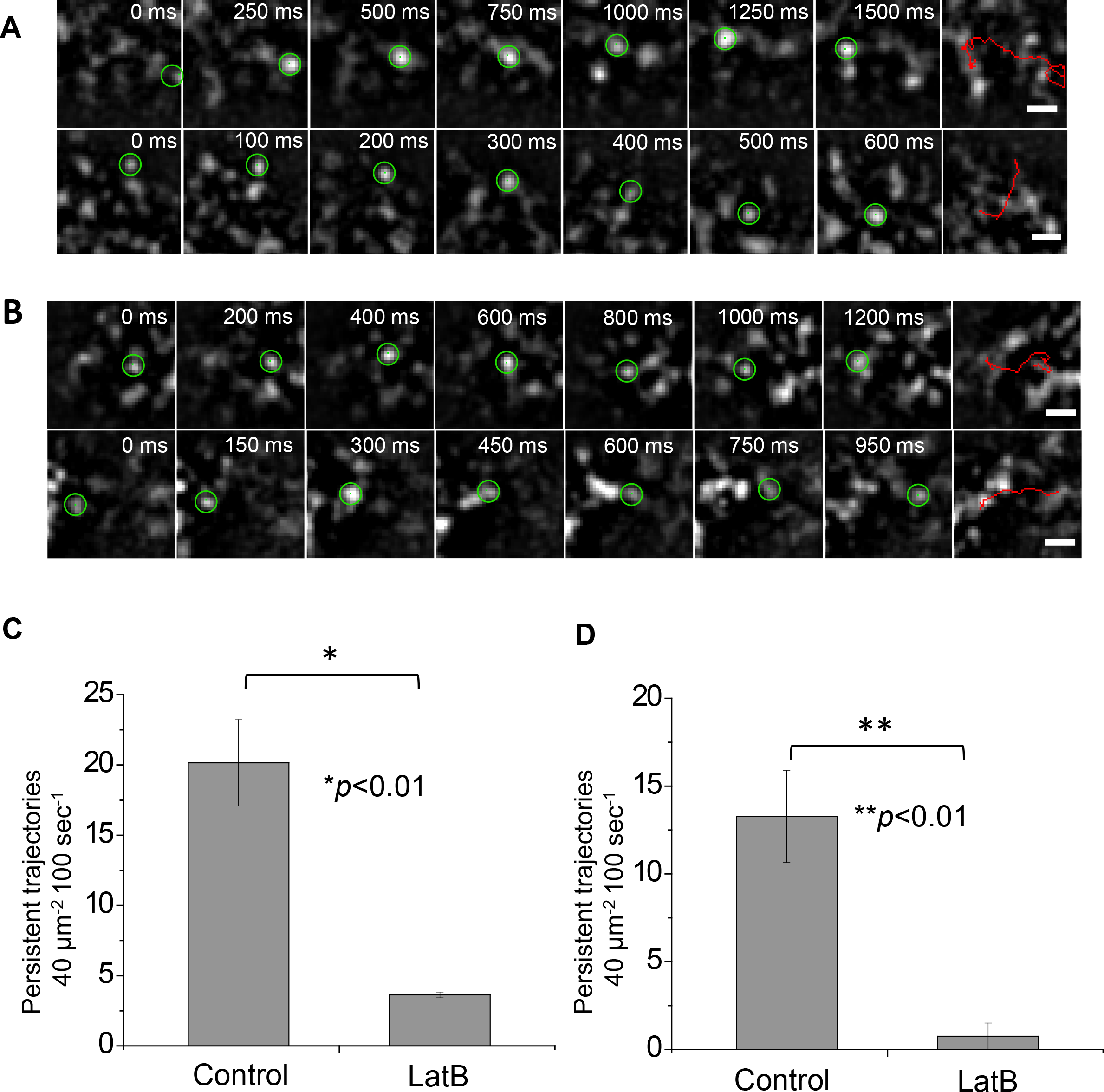
Myosin XIa and VAMP72-labeled vesicle directed motion at the cell cortex depends on actin. **A and B)** Representative examples of persistent linear trajectories of 3mEGFP-Myosin XIa (A) and 3mEGFP-VAMP72-labeled vesicles (B) at the cortex of apical caulonema cells near the apex. Images were acquired using VAEM at 50 ms intervals for the complete time series. Selected images of each trajectory are shown with the time from the series indicated at the top. Persistent linear trajectories are shown in the last image of each series. Scale bar is 1 μm. **C and D)** Quantification of the number of persistent linear trajectories detected in control and latrunculin B treated apical caulonema cells for myosin XIa (n=6 cells for each condition) (C) and VAMP72-labeled vesicles (n=3 cells for each condition) (D). Error bars correspond to the standard errors of the mean.

### Myosin XIa and VAMP72 Labeled Highly Dynamic Punctate Structures at the Cortex of Moss Protoplasts

Although we were able to identify a portion of myosin XIa and VAMP72-labeled vesicles that exhibited actin dependent active motion. To track both myosin XIa and VAMP72-labeled vesicles together, we needed to improve our imaging resolution and tracking fidelity. For this reason, we imaged protoplasts with VAEM; with protoplasts, because it was possible to slightly flatten the cells and reliably get larger imaging areas and more consistent imaging data. Just as in caulonema cells, we found that both fluorescently labeled myosin XIa and VAMP72-labeled vesicles localize to dynamic punctate structures (Figure 4A).

**Figure 4.**
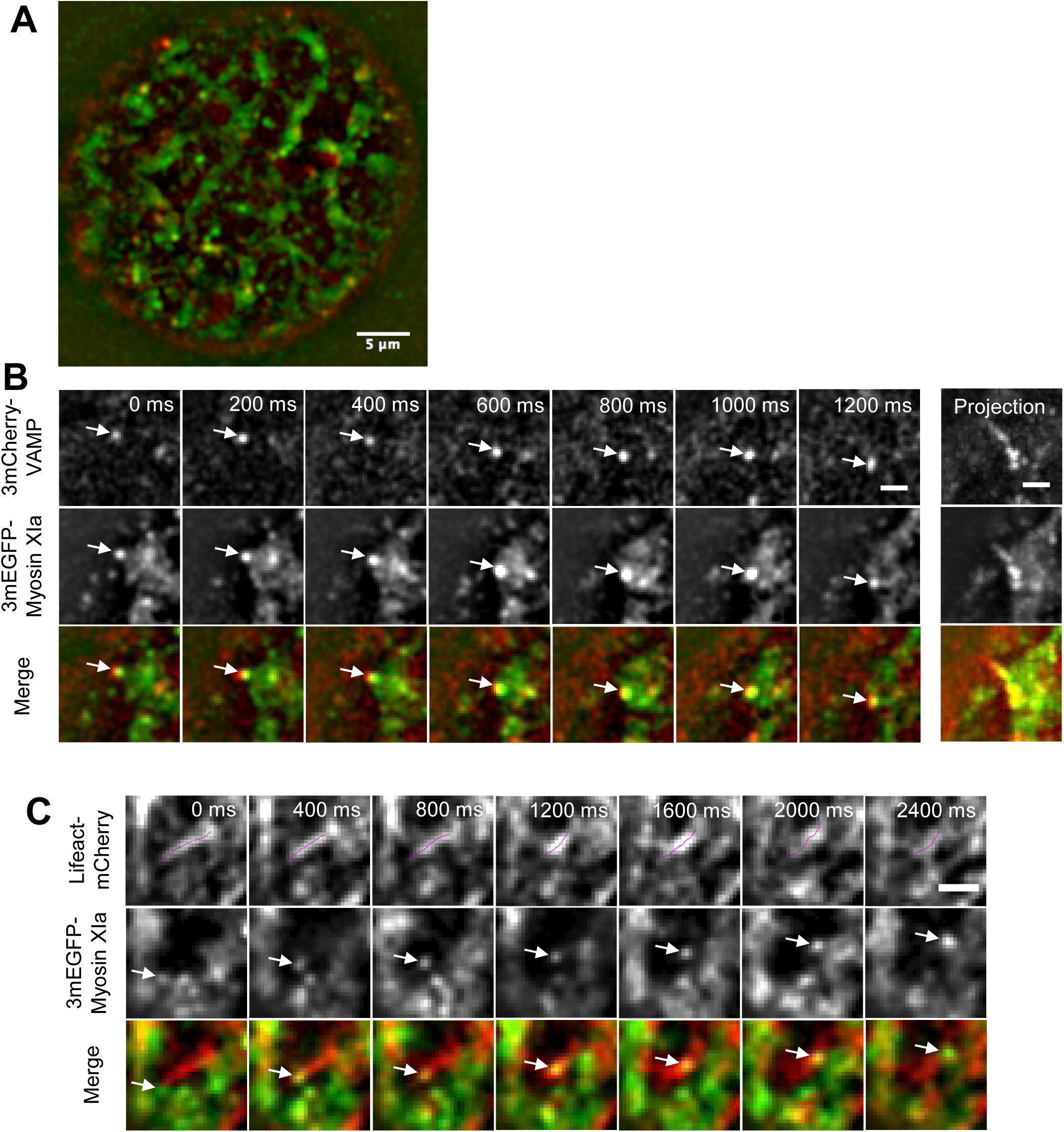
Myosin XIa and VAMP72-labeled vesicles exhibit co-localized dynamic motion in protoplasts. **A)** Representative VAEM image of a protoplast expressing 3mCherry-VAMP and 3mEGFP-myosin XIa. Scale bar 5 μm **B)** Selected images of co-localized 3mCherry-VAMP and 3mEGFP-myosin XIa are shown in the top two rows and the merged image in the third row with VAMP-labeled vesicles in red and myosin XIa in green. Images were simultaneously acquired from protoplasts using VAEM at 100 ms intervals. Maximum projections of 26 frames from each corresponding time series shown in the right panel. Scale bar 2 μm. **C)** Selected images of Lifeact-mCherry and 3mEGFP-myosin XIa are shown in the top two panels and the merged image in the third one with actin filaments in red and myosin XIa in green. Images were simultaneously acquired from protoplasts using VAEM at 100 ms intervals. Arrows show myosin XI-labeled punctuate structures moving on actin filaments depicted by the dotted lines. Scale bar is 2 μm.

In protoplasts, we observed that several punctate structures labeled with both 3mEGFP-myosin XIa and 3mCherry-VAMP72 move in a linear manner, as indicated by the maximum projections in the examples shown in Figure 4B. To test the overlap of these trajectories with actin filaments, we simultaneously imaged 3mEGFP-myosin XIa and Lifeact-mCherry-decorated F-actin (Vidali et al., 2009). We found that myosin XIa-labeled structures co-localize and move along cortical actin filaments in moss protoplasts (Figures 4C). Together with the fact that myosin XIa is an actin-based motor, these results further support that the motion of the endomembrane vesicles labeled by both myosin XIa and VAMP72 occurs on actin filaments.

### Myosin XIa-associated Structures and VAMP72-labeled Vesicles Exhibit Short Persistent Active Motion in Protoplasts

Because of the improved resolution in protoplasts, it was possible to quantitatively infer how myosin XIa and VAMP72-vesicles associate with F-actin by analyzing the persistent nature of their trajectories—without the need for tracking individual actin filaments. To this aim we used a hidden Markov model, (HMM), see Methods and Sup Fig 2 for more details (Rabiner, 1989). Briefly, an HMM is a statistical model for determining the likelihood, that at a given time, a system is in some unobserved state based on a known observable variable. As previously shown (Roding et al., 2014), an HMM can be applied to intracellular particle tracking data to determine the likelihood that, at a specific point in a given trajectory, a particle is either in a Brownian state or an active transport state (Sup Figure 2A). Importantly, this model does not require fluorescently labeling the actin cytoskeleton or tracking individual actin filaments in crowded environments. It only requires the tracking of particles from one channel data (in our case, myosin XIa or VAMP72), which could be reliably done with the Fiji plugin, Trackmate (see Methods).

When the HMM was applied to myosin XIa and VAMP72-labeled vesicles tracking data in protoplasts, it found 139 out of 307 (45 %), and 126 out of 572 (22%), trajectories with an active component, respectively (Figure 5A, 5B and Sup Movies 4-6). The HMM also estimated the most likely state transition probabilities for myosin XIa and VAMP72-labeled vesicles, Table 2 (see Methods). We found that both myosin XIa and VAMP72-labeled vesicles have approximately a 75% chance of remaining on a filament (*P*(*h*_*t*_ = *A*|*h*_*t*−1_ = *A*)) during the 50 ms exposure time. Consistent with this rate, we did not observe any continuously active sections of trajectories longer than 1.05 s. We also observed the actin-associated run lengths to have an average end-to-end distance of 1.24 ± 0.06 and 1.30 ± 0.07 μm for myosin XIa and VAMP72-labeled vesicles, respectively (errors indicate standard error of the mean). The instantaneous velocity for active trajectories of myosin XI and VAMP72-labeled vesicles was 2.7 ± 1.95 and 2.45 ± 1.89 μm/sec, respectively (errors indicate standard deviation). Interestingly, the instantaneous velocity for Brownian trajectories was on a similar scale, with 2.4 ± 2 and 1.83 ± 1.66 μm/sec, for myosin XI and VAMP72-labeled vesicles, respectively (errors indicate standard deviation).

**Figure 5.**
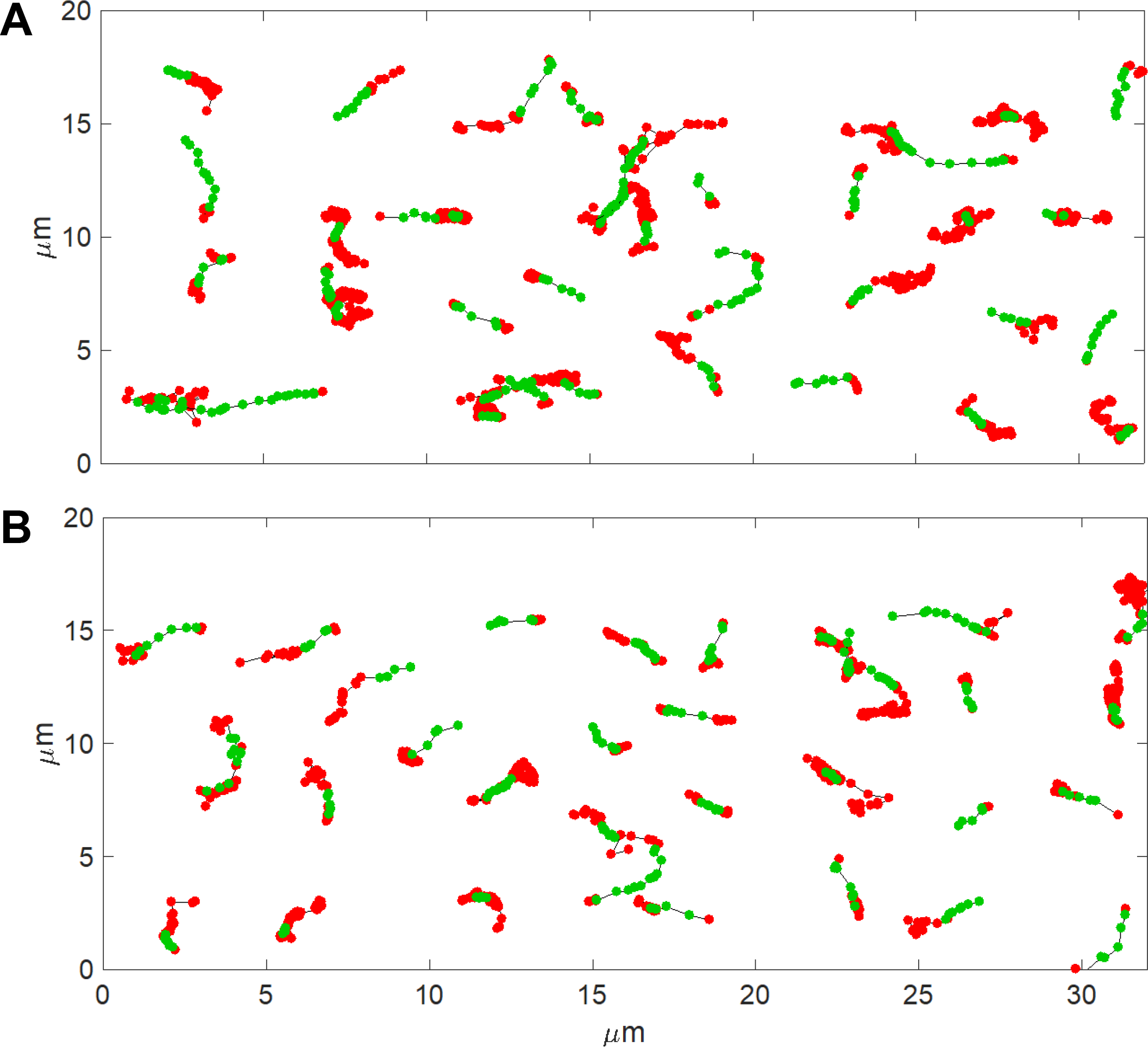
Example hidden Markov model predicted trajectories in protoplasts. **A)** Example VAMP72-labeled vesicle trajectories predicted by the HMM containing a sequence in the active state. **B)** Example myosin XIa trajectories predicted by the HMM containing a sequence in the active state. Red circles indicate the Brownian state and Green circles indicate the active state.

**Table 1.**
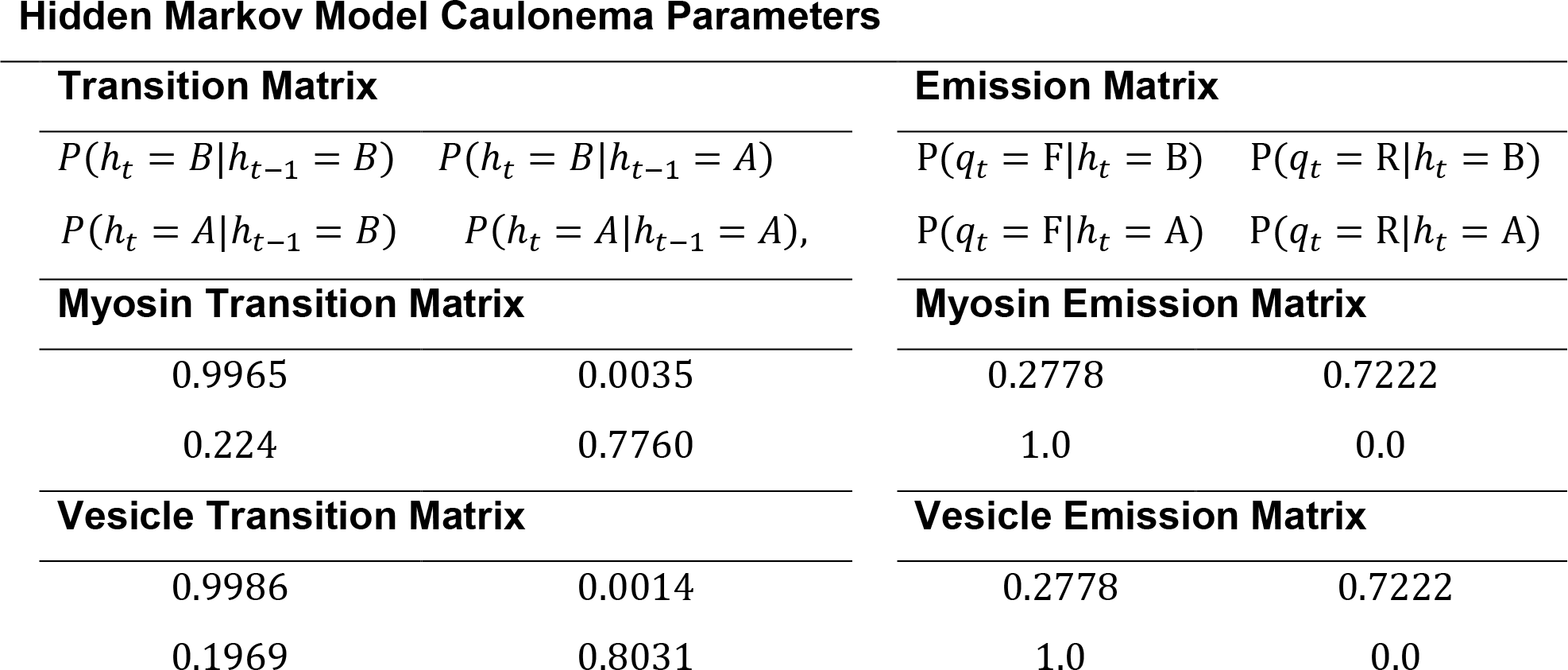
Hidden Markov Model Caulonema Parameters. Transition and emission matrices for 3mEGFP-myosin XIa and 3mEGFP-VAMP72-labeled vesicles measured at the cell cortex. Emission matrices were held constant throughout parameters convergence.

| Transition Matrix |  | Emission Matrix |  |
| --- | --- | --- | --- |
| $P(h_t = B\vert h_{t-1} = B)$ | $P(h_t = B\vert h_{t-1} = A)$ | $P(q_t = F\vert h_t = B)$ | $P(q_t = R\vert h_t = B)$ |
| $P(h_t = A\vert h_{t-1} = B)$ | $P(h_t = A\vert h_{t-1} = A)$ | $P(q_t = F\vert h_t = A)$ | $P(q_t = R\vert h_t = A)$ |
| Myosin Transition Matrix |  | Myosin Emission Matrix |  |
| 0.9965 | 0.0035 | 0.2778 | 0.7222 |
| 0.224 | 0.7760 | 1.0 | 0.0 |
| Vesicle Transition Matrix |  | Vesicle Emission Matrix |  |
| 0.9986 | 0.0014 | 0.2778 | 0.7222 |
| 0.1969 | 0.8031 | 1.0 | 0.0 |

**Table 2.** Hidden Markov Model Protoplast Parameters. Transition and emission matrices for 3mEGFP-myosin XIa and 3mEGFP-VAMP72-labeled vesicles measured at the cell cortex. Emission matrices were held constant throughout parameter convergence. Here A and B indicate the hidden active and Brownian states, and F and R indicate the observed forward and reverse emissions.

| Transition Matrix |  | Emission Matrix |  |
| --- | --- | --- | --- |
| $P(h_t = B\vert h_{t-1} = B)$ | $P(h_t = B\vert h_{t-1} = A)$ | $P(q_t = F\vert h_t = B)$ | $P(q_t = R\vert h_t = B)$ |
| $P(h_t = A\vert h_{t-1} = B)$ | $P(h_t = A\vert h_{t-1} = A)$ | $P(q_t = F\vert h_t = A)$ | $P(q_t = R\vert h_t = A)$ |
| Myosin Transition Matrix |  | Myosin Emission Matrix |  |
| 0.9767 | 0.0233 | 0.2778 | 0.7222 |
| 0.2794 | 0.7206 | 1.0 | 0.0 |
| Vesicle Transition Matrix |  | Vesicle Emission Matrix |  |
| 0.9656 | 0.0344 | 0.2778 | 0.7222 |
| 0.2486 | 0.7514 | 1.0 | 0.0 |

### Myosin XIa and VAMP72-labeled Vesicles Exhibit a Weak and Transient Association

To determine if myosin XIa and VAMP72-labeled vesicles are associated with the same membranes, we simultaneously imaged myosin XIa-labeled structures and VAMP72-labeled vesicles in protoplasts. Using an approach similar to Deschout et al. (Deschout et al., 2013), we used automated analysis to determine if the myosin XIa and VAMP72-labeled vesicles exhibited spatiotemporal co-localization. In this approach, we used a minimal contact radius and a sliding correlation window to determine if two particles from different channels showed correlated movement (see Methods and Sup Figure 3). For each given window size, the frequency of VAMP72-labeled trajectories was calculated by dividing number of VAMP72-labeled trajectories co-localizing with a myosin XI, by all the VAMP72-labeled trajectories (Figure 6A and Methods). We found that a fraction of myosin XIa-decorated structures partially co-localize —in time and space— with VAMP72-labeled vesicles at the cell cortex in moss protoplasts. We observed that at least 20% of the VAMP72-labeled trajectories exhibited some co-localization during a total period of 400 ms, and at least 15 % during a total period of 1 s (Figure 6A and Sup Movies 6-7). These co-localization frequencies are significantly higher than those obtained by randomly shuffling the observed trajectories (Figure 6A, Random Shuffle). The reduction in localization frequency at increasing window sizes is the result of increased stringency for co-localization at bigger window sizes.

**Figure 6.**
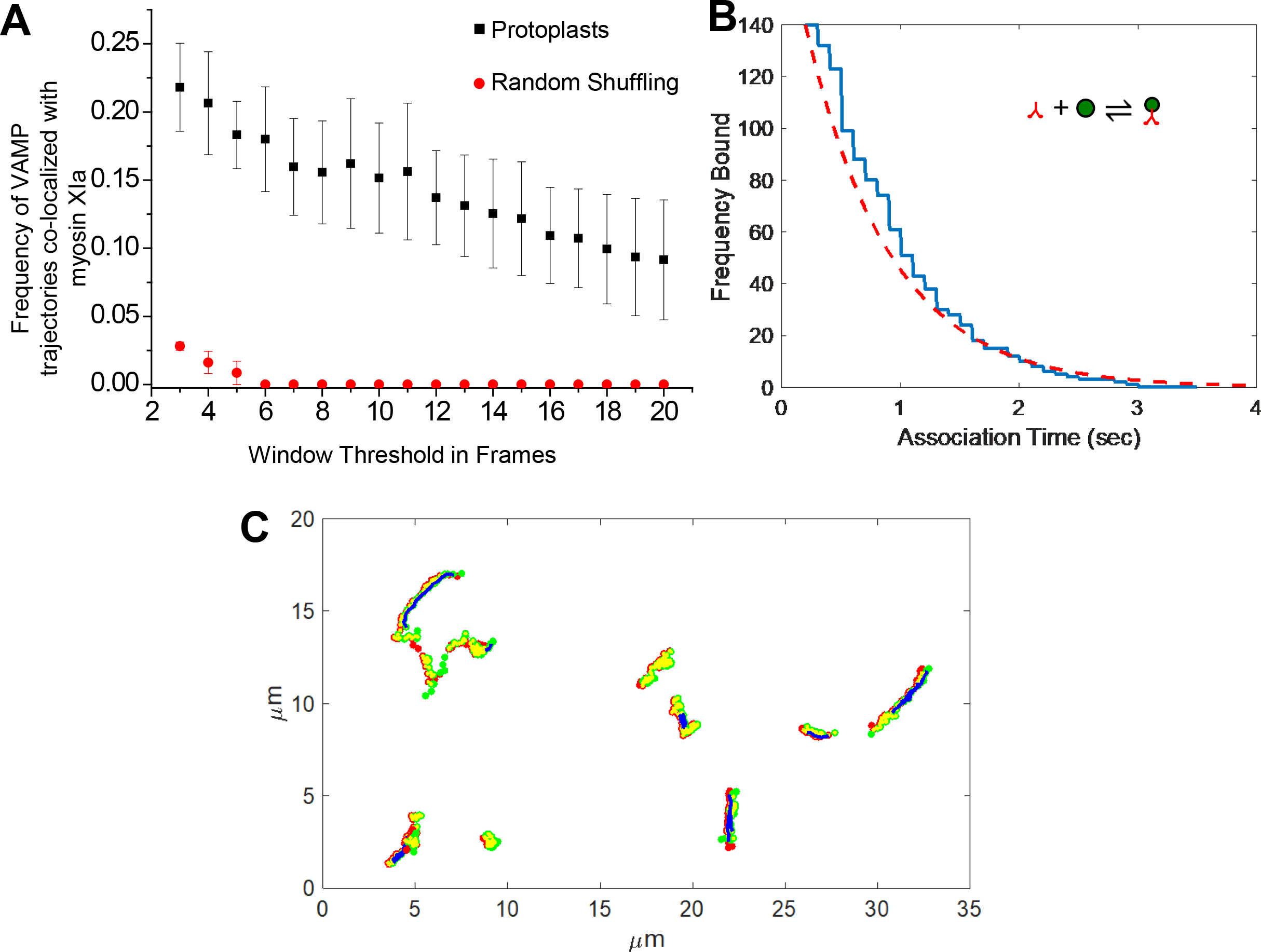
Myosin XIa colocalization with VAMP72-vesicles is specific and transient. **A)** Frequency of 3mCherry-VAMP72 trajectories that contain co-localization with 3mEGFP-myosin XIa detected in protoplasts (black square) or randomly generated by a simulation (red circle). x-axis represents the minimum required number of frames two particles have to be moving together to be classified as co-localized. n=3 cells. **B)** The number of myosin XIs observed on a vesicle in protoplasts for a given time (blue line) and the corresponding best fit to Eq. 7 (red dotted line). **C)** Example co-localized protoplast trajectories. VAMP72-labeled vesicle trajectories (red dots) and myosin XIa trajectories (green dots) co-localized (yellow dots) exhibiting active motion (blue lines) predicted by the HMM.

To examine the total co-localization time of both myosin XIa and VAMP72-labeled vesicles, we divided the total number of co-localization events by the total number of identified events for a window size of 3 frames. Of all identified and tracked myosin XIa, we found that ~4.2% of it was localized to vesicles; we also found that 10% of the identified VAMP72-labeled vesicles were co-localized with a myosin XI. Although these percentages may seem low when compared with our FRAP results, they result from a co-localization analysis that requires spatial co-localization specifically at the cell’s cortex. Nevertheless, because of the stringency of this analysis, these measurements are well above random co-localization events (Figure 6A). This result is consistent with the hypothesis that myosin XIa associates with endomembrane vesicles.

The co-localization data also allow us to estimate the off-rate for the myosin XI-VAMP72-labeled vesicle interaction. To determine this rate, we examined the duration myosin XIa resided on a vesicle. Fitting the observed residence times to an exponential decay of the form, i.e.,

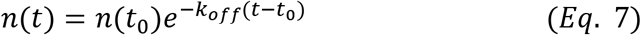

yields an off rate of *k_off_* ~ 1.4 s^−1^ (Figure 6B). Here *n* is the number of molecules bound after the elapsed time t, and t_0_ is the minimum imaging required to detect a co-localization event (200 ms) by the co-localization algorithm. This off-rate is fast, resulting in a half-life for the complex of ~500 ms.

To determine if VAMP72-labeled vesicle motion is dependent on myosin XIa we examined the HMM results and the co-localized trajectories together, Figure 6C and Sup Movies 4-7. We found that of the 139 VAMP72-labeled vesicle active trajectories only 28(20%) of them were co-localized with a detectable myosin XIa, Sup Movie 6. Since 80 % percent of the VAMP72-labeled vesicles were able to move actively without a myosin XIa (Sup Movie 4), this is likely due to a population of VAMP72-labled vesicles propelled by formin-mediated actin polymerization (Furt et al., 2013; van Gisbergen et al., 2012). Since 19 out of the 126 (15 %) myosin XIa active trajectories were co-localized with a detectable VAMP72-labeled vesicle (Sup Movie 6), myosin XIa may be able to move on actin without any cargo or myosin XIa might transport another cargo other than VAMP72-labeled vesicles (Sup Movie 5).

## Discussion

In this work, we have shown for the first time to the best of our knowledge, that myosin XIa and VAMP72-labeled vesicles co-localize *in vivo*, and exhibit actin dependent mobility that was observed as short persistent trajectories. Using particle-tracking statistics, we found that myosin XIa exhibits a transient interaction with VAMP72-labeled vesicles with a fast off-rate (~ 1.4 s^−1^). Taken together, our results support a hypothesis in which myosin XIa transiently interacts with its binding constituents to flexibly coordinate vesicle motion and actin dynamics.

To better understand how myosin XIa and VAMP72-labeled vesicles interact with the actin cytoskeleton at the tip of the cell during polarized growth we first performed Fluorescence Recovery After Photobleaching experiments (FRAP). Here we found that myosin XIa and VAMP72-labeled vesicles exhibited an actin-dependent fluorescence recovery. During fluorescence recovery, we found that both myosin XIa and VAMP72-labeled vesicles restored the apical localization at the cell tip in under 2 s. Considering that we used a bleach radius of 2 μm and that the myosin XIa and VAMP72-labeled vesicles exhibit transport velocities around 2 μm/s, these recovery rates are consistent with active transport. However, we did find that myosin XIa was able to achieve apical localization before VAMP72-labeled vesicles. This indicates that myosin XIa displays an additional tip-localized recovery mode different from simple diffusion or VAMP72 vesicle association. This recovery mode maybe unloaded myosin XIa moving on actin or myosin XIa exchanging off of bleached cargo at the tip, identifying the mechanism for this additional recovery is left for future work. Nevertheless, the apical recovery rates seen here, and the myosin XIa movement without VAMP72-labeled vesicles, are consistent with the active transport found by the HMM at the cell cortex

To demonstrate that myosin XIa associates with vesicles, we performed Variable Angle Epifluorescence Microscopy (VAEM) on protoplasts expressing fluorescently labeled myosin XIa and VAMP72-labeled vesicles. We found that myosin XIa co-localizes with VAMP72-labeled vesicles for periods consistent with a an off-rate of 1.4 s^−1^. This off-rate is an upper limit estimate because we cannot be certain when each association occurred during our imaging time. Additionally, we cannot be certain when a particle leaves or diffuses out of the evanescent field. Nevertheless, this suggest that myosin XIa and VAMP72-labeled vesicles do not reside together much longer than a few seconds. This is also supported by our FRAP analysis, where a fraction of 55%-66% of myosin is associated with vesicles. In contrast, using single particle tracking, we found that only a small fraction of the tracked myosin XIa (4.3%) was associated with VAMP72-labeled vesicles. This discrepancy is likely due to limitations of the VAEM system, which relies on a narrow evanescent field, low signal to noise ratio, fluorophore bleaching, and relatively slow acquisition rate. The important conclusion is that a significant fraction of myosin XIa molecules remain free. Hence, our measurements suggest that *in vivo,* there is a significant population of myosin XIa and vesicles that exhibit brief transient interactions.

We also observed for the first time, to the best of our knowledge, several instances in which fluorescently labeled myosin XIa and VAMP72-labeled vesicles associated and moved along Lifeact decorated actin filaments. Using a Hidden Markov Model (HMM) on particle tracking data we were able to characterize active motion. In both caulonema and protoplasts, we found that both myosin XIa and VAMP72-labeled vesicles moved in an active manner for distances not much greater than a few microns for periods of less than a second. These trajectories were completely abolished in caulonema treated with latrunculin B, indicating that these active trajectories are actin dependent.

To better understand how the interaction between myosin XIa and VAMP72-labeled vesicles influences their active motion, we combined our co-localization and HMM analyses. We found that active motion of myosin XIa and VAMP72-labeled vesicles could happen with or without an observed co-localization. This indicates that myosin XIa either transports cargo other than VAMP72-labeled vesicles or that myosin XIa can move unloaded on filamentous actin. Unloaded myosin XIa on F-actin could be important for filament organization and maintenance. VAMP72-labeled vesicles may also move through the formin-mediated actin propulsion system previously suggested {Schuh, 2011 #1095; Furt, 2013 #1199}. We do not believe that there is any microtubule dependent motion of VAMP72 because latrunculin abolished active motion in caulonema. Although we observed no localization to myosin XI in some cases, it is possible that this VAMP72-labeled active motion is myosin XI dependent and we do not have the resolution to detect it.

Based on the particle tracking evidence and the bulk FRAP measurements, we conclude myosin XIa and VAMP72-labeled vesicles exhibit a transient interaction between each other and exhibit short run lengths on actin filaments. Since *P. patens* does not exhibit large organelle cytoplasmic streaming (Furt et al., 2012), it is not surprising that myosin XIa only displays short-lived transport of vesicles. Additionally, the myosin XIa actin-transport speeds found here are slightly slower relative to other orthologs (Tominaga et al., 2013) and maybe a product of the slower polarized growth found in moss. Importantly, the weak interactions and short run lengths we observed are consistent with a mechanism by which myosin XIa can actively remodel and regulate the oscillating apical F-actin, during polarized growth, while still promoting vesicle focusing for exocytosis see Figure 7. Without the flexibility of these weak transient interactions, the apical actin spot may not be as dynamic or easily regulated. Furthermore, the quantitative exploration of these binding interactions can begin to help shape future modeling studies that incorporate the cytoskeleton in polarized cell growth.

**Figure 7.**
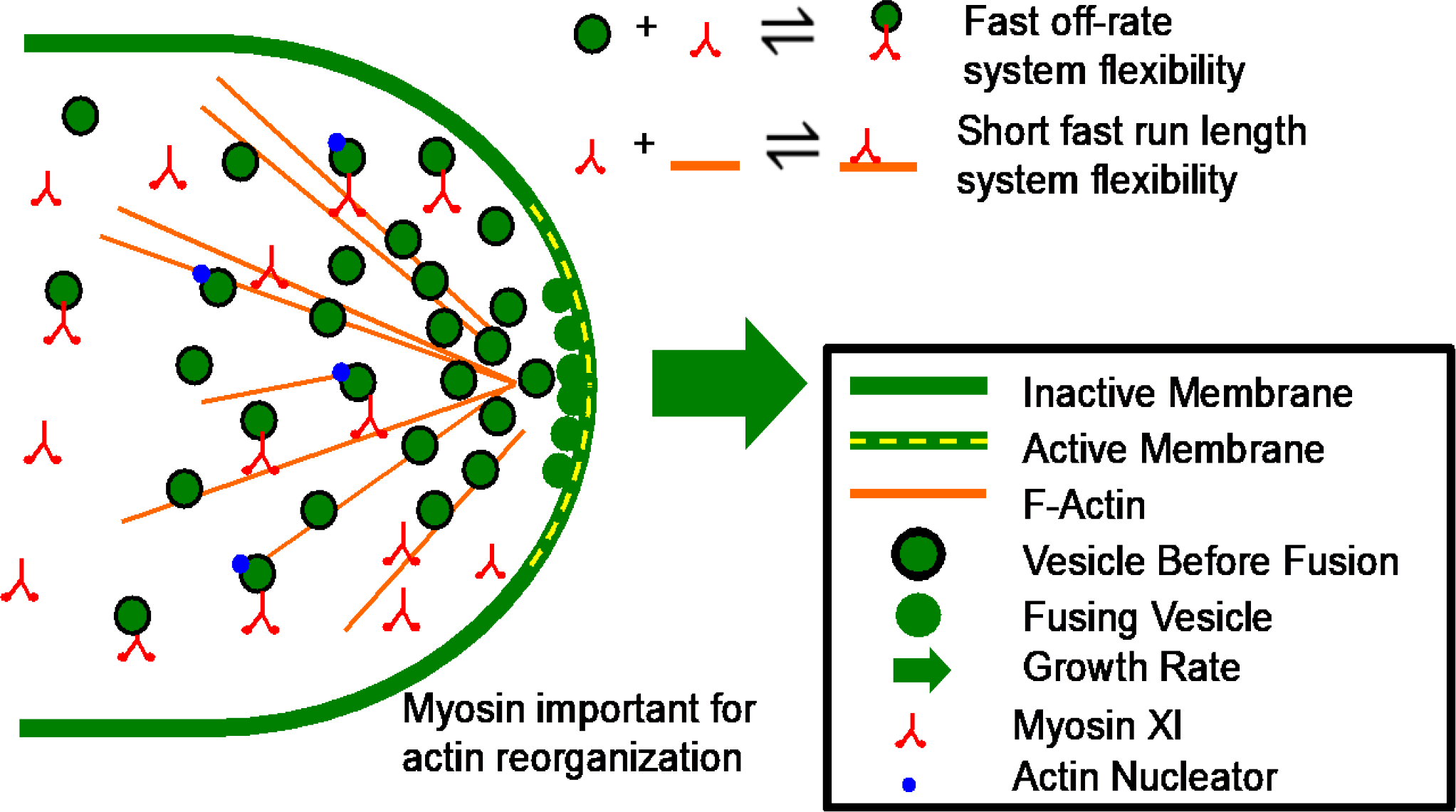
Transient interactions between myosin XIa, VAMP72-labeled vesicles, and F-actin give rise to system flexibility in polarized cell growth. Here myosin XIa and vesicles transiently interact with each other and the actin cytoskeleton to give rise to short lived persistent motion directed toward the secretion zone at the cell apex. These transient interactions allow for the dynamic actin oscillations observed during tip growth while at the same time providing adequate vesicle focusing to sustain wall expansion.

## Methods

### Constructs and Cell Lines

Two independent 3mEGFP-VAMP72 lines were obtained by transforming the Gransden strain with pTH-Ubi-3mEGFP-VAMP72, and by selecting for stable lines. The construct was obtained by a two element LR Gateway reaction of entry clones pENT-L1-3mEGFP-R5 and pENT-L5-VAMP72-L2 (Furt et al., 2013) and the destination vector pTH-Ubi-Gateway (Vidali et al., 2007). The 3mCherry-VAMP72 plus 3mEGFP-MyosinXIa and the Lifeact-mCherry plus 3mEGFP-MyosinXIa lines were previously described (Furt et al., 2013).

### Culture Conditions

*Physcomitrella* cell lines were propagated using standard methods as previously described (Vidali et al., 2007). Isolation of protoplasts and DNA transformation were done according to Liu and Vidali (Liu and Vidali, 2011).

For the VAEM imaging (Konopka and Bednarek, 2008), moss cell lines were grown on PpNO_3_ medium (Vidali et al., 2007) for 7 days. One week-old protonemal tissue was either directly mounted on glass microscope slide for imaging or used to prepare protoplasts as previously described with some modifications (Liu and Vidali, 2011). After counting, protoplasts were resuspended in 5 ml of 8 % (w/v) mannitol and cultured in the growth chamber for one day. For imaging, moss protonemal tissue or protoplasts were deposited on a 1 % (w/v) agar pad in PpNO_3_ medium on a glass microscope slide. 10 μL of liquid PpNO_3_ medium was applied before covering with a glass 0.25 mm thick coverslip and sealing with VALAP (1: 1: 1, VAseline, LAnolin, Paraffin). For latrunculin B treatments, 10 μL of 25 μM latrunculin B resuspended in PpNO_3_ medium was added to the preparation instead of the PpNO_3_ medium alone. Apical caulonemal cells were imaged exactly 10 min after the treatment. For FRAP, moss cell lines were grown and cultured as described in (Bibeau et al., 2018).

### VAEM Microscopy

Protoplasts were mounted on an inverted microscope (model Ti-E; Nikon Instruments) and imaged with an Apo TIRF, 60 ×, NA 1.49, oil immersion objective (Nikon Instruments). Apica caulonemal cells from protonemal tissue were imaged with an Apo TIRF, 100x, NA 1.49, oil immersion objective (Nikon Instruments). To increase magnification, the 1.5x optivar was used to collect all images. The 488 and 561 nm laser lines were used to excite mEGFP and mCherry, respectively. To provide the maximum signal-to-noise ratio, the laser illumination angle was finely tuned for each sample. Signals were simultaneously acquired for both channel using a beam splitter and a 512×512 electron-multiplying CCD camera iXON3 (Andor Technology).

### Image Processing and Tracking

Images collected by VAEM, were first processed in ImageJ using a background subtraction (rolling ball radius was tuned to 5 for myosinXI and VAMP72), a FFT bandpass filter (a diameter comprised between 2 and 20 pixels was used) and an enhance contrast tool (enhancement with no pixel saturation was normalized to all slices of each time series). Detection and tracking of the cortical punctate structures of myosin XIa and VAMP72 were performed using the TrackMate plugin in FIJI. We manually tracked several punctate structures to determine the best settings. Briefly, the DoG Detector tool was chosen because it is optimal for small spot sizes and it allows to differentiate between two spots that are close to each other. The diameter of the structures was no greater than 0.6 μm and the threshold was kept at 1000 for all molecules analyzed. For tracking, we used the simple LAP tracker tool with the following parameters: linking maximum distance of 0.5 *μm*; gap-closing maximum distance of 0.7 μm; gap-closing maximum frame gap of 0. Two other filters were applied to track the punctate structures that move in a linear persistent manner: a number spots per track greater than 12 (equivalent to 600 ms) and a displacement greater than 1.2 μm. Each trajectory recorded was verified manually using an in-lab developed Matlab routine. To track the punctate structures that stay confined to an area, we adjusted the filters as follow: a number of spots per track greater than 90 (equivalent to 4.5 s) and a displacement no greater than 0.5 μm. The number of trajectories was normalized by average imaged area (40 μm^2^) for each cell. The instantaneous velocities of each tracked punctate structure were calculated using the coordinates obtained by the simple LAP tracker tool.

### Co-localization Frequency

To automate the identification of co-localized fluorescently labeled myosin XIa and VAMP72-labeled vesicles we used an approach similar to (Deschout et al., 2013). With a custom Matlab (Mathworks Natick, MA) routine, we compared the myosin XIa and VAMP72-labeled trajectories. An example analysis can be found in Sup Figure 3. For two trajectories to be considered co-localized they first needed to have a temporal overlap and be within a specified contact radius of 0.5 μm. Then a sliding correlation window was used to determine the correlations between the two channels in the x and y directions, respectively. If the correlation within the window was greater than 0.7 the movement was classified as correlated for that direction. Window correlations were found using the Matlab function, *corr*. If the trajectories exhibited correlated motion in both the x and y directions the time point within the selected window was classified as co-localized. Here we used a minimum correlation threshold of 0.7 and a maximum contact radius of 0.5 μm to account for tracking and alignment error.

To demonstrate that the observed co-localized trajectories were not a result of random Brownian motion, we randomly shuffled the existing trajectories in time and space with a custom written Matlab script. The new trajectories were then analyzed for co-localization as previously mentioned.

### Hidden Markov Model

To facilitate the use of a Hidden Markov Model (HMM), trajectories were preprocessed such that they were converted from spatial coordinates into either reverse, *R*, or forward, *F*, moves (Roding et al., 2014). Specifically, we measured the angle between three consecutive points along the trajectory, namely, *t*_*i*−1_, *t*_*i*_, and *t*_*i*+1_. This angle *θ*_*i*_ was then classified as a forward move if the angle was greater than a specific threshould or classified as a reverse. Trajectories were also filtered by size such that, trajectories shorter than 7 frames or longer than 150 frames were not used.

To classify the transport state of a particle for a given point on these modified trajectories, we employed a Hidden Markov Model (HMM) (Roding et al., 2014)(Sup Figure 2) with two hidden states, namely, an active state, *A*, and a Brownian state, *B*. Essentially, the HMM determines the number of consecutive forward moves Here, we represent each *hidden* sequence as *H*=*h*_*1*_,*h*_*2*_,…,*h*_*T*_ and each *observed* sequence as *Q*=*q*_*1*_,*q*_*2*_,…,*q*_*T*_. In the active state, *A*, we assumed that myosin XIa and VAMP72-labeled vesicles always move forward, while in the Brownian state, *B*, we assumed an equal probability distribution over all angles, *θ*. Thus the emission probabilities in the Brownian state are only a function of the angle threshold, where the probability of moving forward given the Brownian state and the probability of moving in the reverse direction given the Brownian state are, *P*(*q*_*t*_ = *F* | *h*_*t*_ = *B*) = (180-130)/(180) and *P*(*q*_*t*_ = *R* | *h*_*t*_ = *B*) = 1 - *P*(*q_t_* = *F* | *h_t_* = *B*), respectively. This leaves the state transition probabilities as the only open parameters in our model. The state transitions probabilities are the chances that protein of interest will change its state on each successive portion of the sequence. The four possible state transitions are as follows, active to active, active to Brownian, Brownian to Brownian, and Brownian to active which can be written formally as *P*(*h_t_* = *A* | *h_t_*−1 = *A*), *P*(*h_t_* = *A* | *h_t_*-1 = *B*), *P*(*h_t_* = *B* | *h_t_*-1 = *B*), and *P*(*h_t_* = *B* | *h_t_*-1 = *A*), respectively.

To find the most likely state transitions for each experimental condition, we used the forward and reverse trajectories as inputs to a modified version of the Matlab function *hmmtrain* for known emission matrices. *hmmtrain* uses the Baum Welch algorithm (Rabiner, 1989), which recursively finds the transition and emission matrices that maximize the likelihood of the observed sequence. We modified this algorithm such that the emission matrix did not change. This modified function reliably found the transition matrix that maximized the likelihood of the observed sequence *Q*, regardless of the initial guess for the transition matrix (Sup Figure 4A and 4B). With the most likely state transitions and the converted sequences, we then used the Matlab function *hmmviterbi* to determine the most likely state of a specific particle at a specific time.

Here we used an angle threshold of 130° was chosen because it allowed for the detection of shorter active trajectories while minimally effecting the most likely transition probabilities (Sup Figure 4C and 4D). Larger angles placed too stringent of a requirement on forward moves and began to influence the transition matrix (Sup Figure 4C and 4D).

### Experimental FRAP Processing

Experimental FRAP was processed as previously described in, (Bibeau et al., 2018). Briefly, FRAP confocal images were saved as tiff stacks and converted into intensity traces by averaging the mean fluorescence intensity within the *4 μm* in diameter region of interest, ROI, using a custom written imageJ macro. All replicate experiments were then averaged and normalized by acquisition photobleaching controls with Matlab. To quantify the spatial fluorescence recovery FRAP tiff stacks had the respective ROIs cropped and averaged using a custom written imageJ macro. Then the fluorescence intensities of the perimeters of the cropped ROIs were extracted, limited volume corrected, and plotted in Matlab.

### Estimating the Myosin XI Diffusion Coefficient

To estimate the myosin XI diffusion coefficient, we used the sedimentation coefficients of myosin V (Krementsov et al., 2004; Li et al., 2004; Wang et al., 2004). Sedimentation coefficients are used to quantify how quickly a protein of interests moves through gradients during ultracentrifugation. Using force balance analysis, it is possible to extract the hydrodynamic radius of a protein based on its sedimentation coefficient (Erickson, 2009), ie,

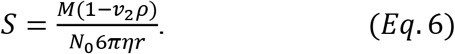

Here *S* is the sedimentation coefficient, which is represented in 10^−13^ seconds, *M* is the molecular weight of the protein of interest, *V*_*2*_ is the inverse density of the protein, *ρ* is the density of the solvent, *N*_*0*_ is Avogadro’s number, *η* is the viscosity of the solvent, and *r* is the hydrodynamic radius of the protein. Based on Ca^2+^ concentrations, it has been shown that myosin V can either be in an open or closed configuration and its sedimentation coefficients were found to be 10.7S and 14S, respectively (Wang et al., 2004) and this value is in agreement between laboratories (Krementsov et al., 2004; Li et al., 2004). Based on *Eq*. 6 and the parameters measured in Wang *et al.*, the corresponding hydrodynamic radii are *13.4* and *10.2 nm* for the open and closed configurations, respectively. Applying the Stokes-Einstein equation for the two configuration’s hydrodynamic radii for myosin V to myosin XI in moss yields theoretical diffusion coefficients of 2.3 and 3.4 μm^2^/s for the open and closed configurations, respectively.

### Modeling Reaction Diffusion FRAP

To solve the reaction diffusion equations Eqs. 1 and 2 in COMSOL (Comsol Inc, Stockholm, Sweden) we used no flux boundary conditions inside a two-dimensional approximation of the moss geometry. We also used initial conditions in which *U(x*,*t*,0) = *total/(1*+*K*_*d*_^*^) and *B(x,y,0)=total-U(x,y,0)* for everywhere outside the 4 *μm* diameter circular bleach region, *R*. Here the effective equilibrium constant is the ratio between the effective on-rate and off-rate, *K*_*d*_^*^*=k*_*on*_^*^/*k*_*off*_, and *total* is the arbitrary combined concentration of the unbound and bound myosin XIa. This concentration is arbitrary, because we observe experimentally the normalized fluorescence intensity within the region of interest, i.e.,

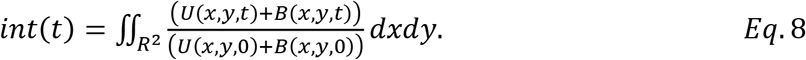

Inside the bleach region the initial concentrations for *U(x,y,0)* and *B(x,y,0)* were multiplied by the bleach depth parameter, *BD*, to simulate an instantaneous ideal bleach.

To solve these equations for arbitrary *k*_*on*_^*^ and *k*_*off*_ we used the finite element modeling software Comsol Multiphysics (Comsol Inc, Stockholm, Sweden). Simulations were created using the “two-dimensional transport of dilute species” interface with two species, a “time-dependent solution”, and a “normal physics controlled mesh”. We then performed a parameter sweep across potential values of *k*_*on*_^*^ and *k*_*off*_ using the Matlab liveLink Comsol Multiphysics module and calculated the sum of squares differences between the simulated recoveries and those measured experimentally. Based on these results we concluded that we could only determine the effective dissociation constant, *K*_*d*_^*^=*k*_*on*_^*^/*k*_*off*_. To converge on a best-fit *K*_*d*_^*^ we imposed gradient descent on the following cost function,

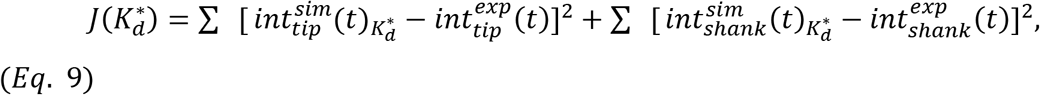

using a custom written script for the Matlab liveLink Comsol Multiphysics module. Here the superscripts *sim* and *exp* indicate simulated and experimental FRAP recoveries, the subscripts indicate the position of the photo-bleach, and *J*(*K*_*d*_^*^) is the dissociation constant dependent cost function. Here we used the numerical forward approximation of a derivative to determine the local gradients during convergence.

## Author Contributions

F.F. performed VAEM experiments, image processing, data analysis, and wrote part of the manuscript. J.B. performed FRAP experiments, developed the HMM model, developed the finite element FRAP model, and wrote part of the manuscript. J.L.K. and I.M. developed DCMS. M.L. helped develop the HMM model. E.T. supervised modeling. L.V. supervised experiments and modeling and edited the manuscript.

## Acknowledgements

We thank all members of the Vidali and Tuzel labs for helpful discussions and to Prof. Arne Gericke for allowing us to use his TIRF microscope. This work was supported by National Science Foundation CBET 1309933 to ET, and NSF-MCB 1253444 to LV. ET and LV also acknowledge support from Worcester Polytechnic Institute Startup Funds.

---

**Sup Figure S1.**
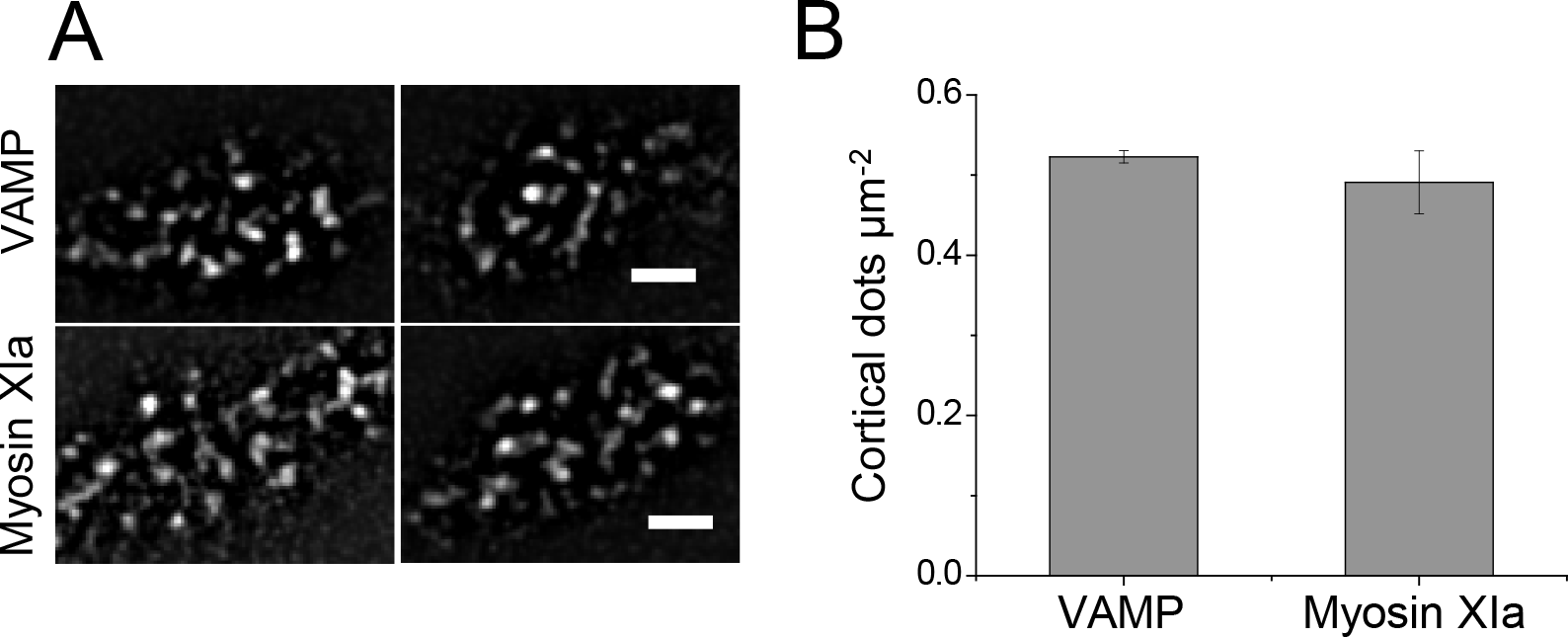
VAMP72 and Myosin XIa localize to punctate structures at the cell cortex of moss cells. **(A)** Two representative images of cortical localization of 3mEGFP-VAMP72 and 3mEGFP-Myosin XIa in apical caulonemal cells acquired using VAEM. Scale bars are 2 µm. **(B)** Average density of cortical punctate structures of 3mEGFP-VAMP72 and 3mEGFP-Myosin XIa in apical caulonemal cells. n=6 cells for 3mEGFP-Myosin XIa and n=3 cells for 3mEGFP-VAMP72. Error bars correspond to the standard errors of the mean.

**Sup Figure S2.**
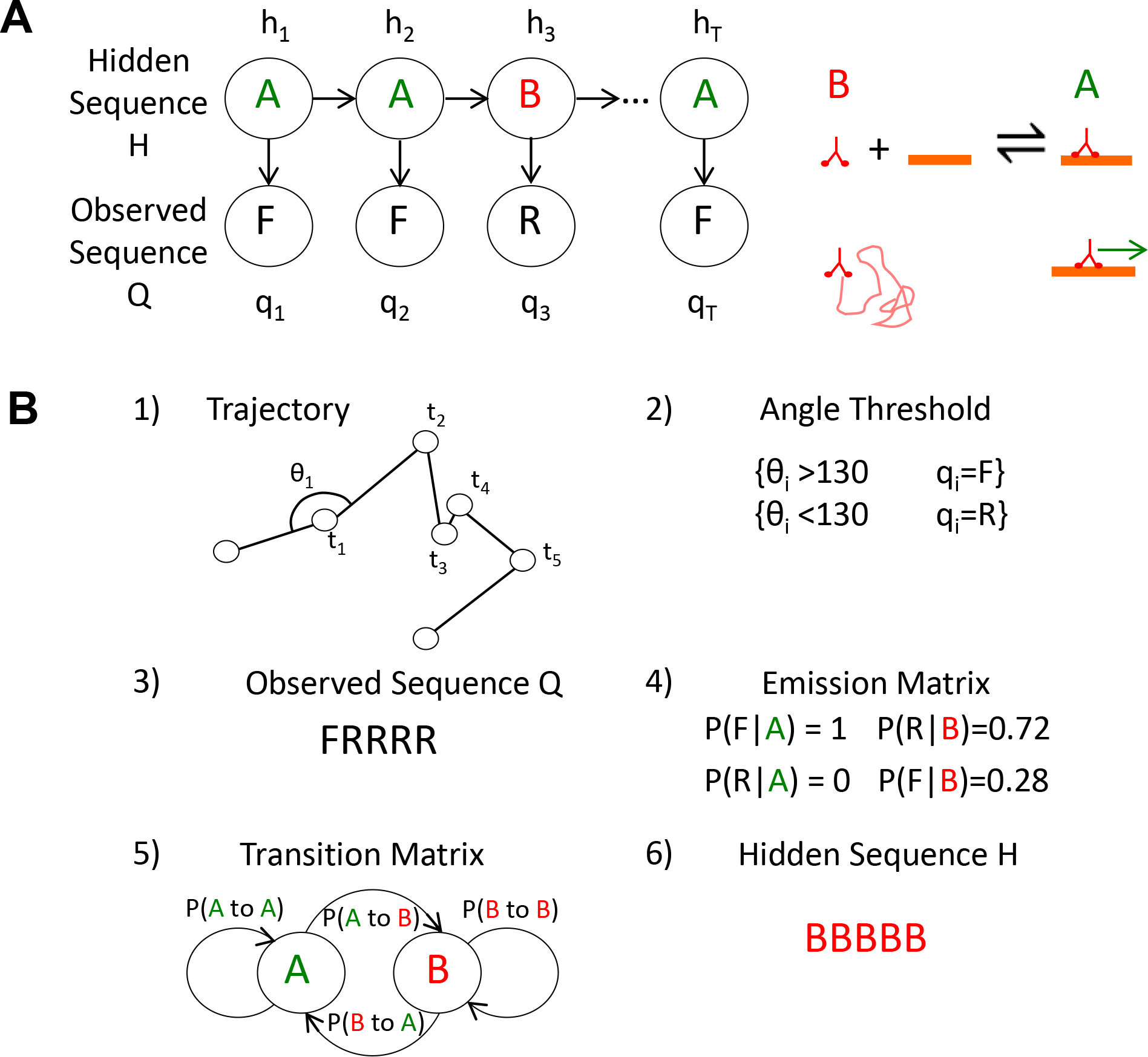
Hidden Markov model explained. A) Example hidden sequence *H* and the corresponding observed sequence *Q*. In the hidden sequence green *A*s correspond to the active transport state and red *Bs* correspond to the Brownian state. In the observed sequence, black *Fs* correspond to forward moves and black *Rs* correspond to backward moves. B) Example of trajectory conversion from observed sequence *Q* to the hidden sequence *H*. Step 1) indicates how the angle θ_i_ was measured across the trajectory. Step 2) shows how that angle was thresholded at 130° to produce forward and backward moves. Step 3) shows the final converted observed sequence *Q*. Step 4) Assume given emission matrix. Step 5) Find transition matrix that maximizes the likelihood of the observed sequence *Q*. Step 6) Find most likely hidden sequence *H*.

**Sup Figure S3.**
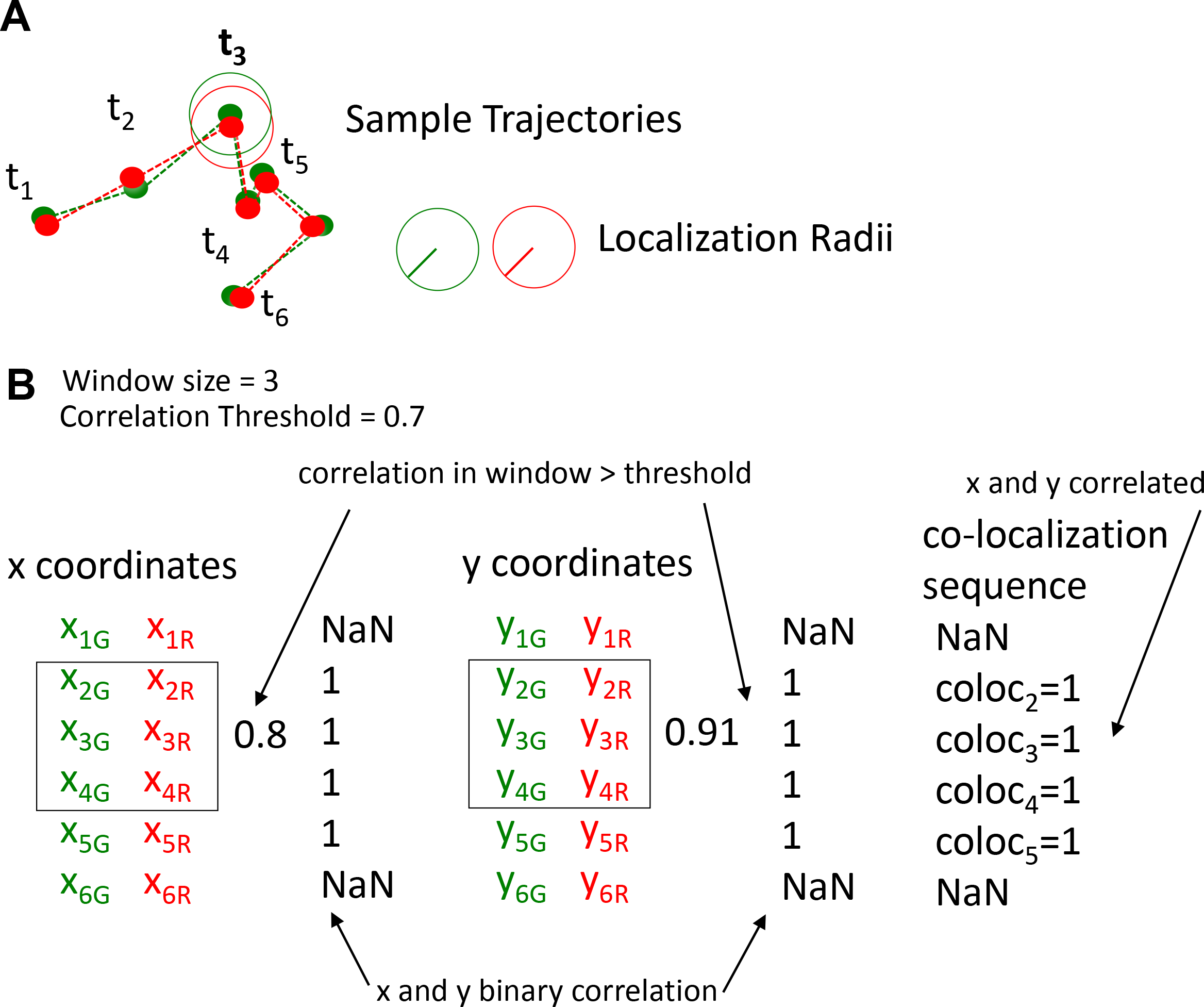
Example co-localization analysis. A) Sample trajectories from the red and green channels and the specified localization radius for t_3_. B) Correlation analysis for x and y coordinates at t_3_ for a window size of 3 frames. Coordinate colors indicate channel. In the example the red and green channels exhibit a correlation above the given threshold in both the x and y directions and are classified as 1. Because both directions are correlated, t_3_ is classified as co-localized. coloc_2_ and coloc_6_ are labeled NaN because the window size does not allow for co-localization at these points in the trajectory.

**Sup Figure S4.**
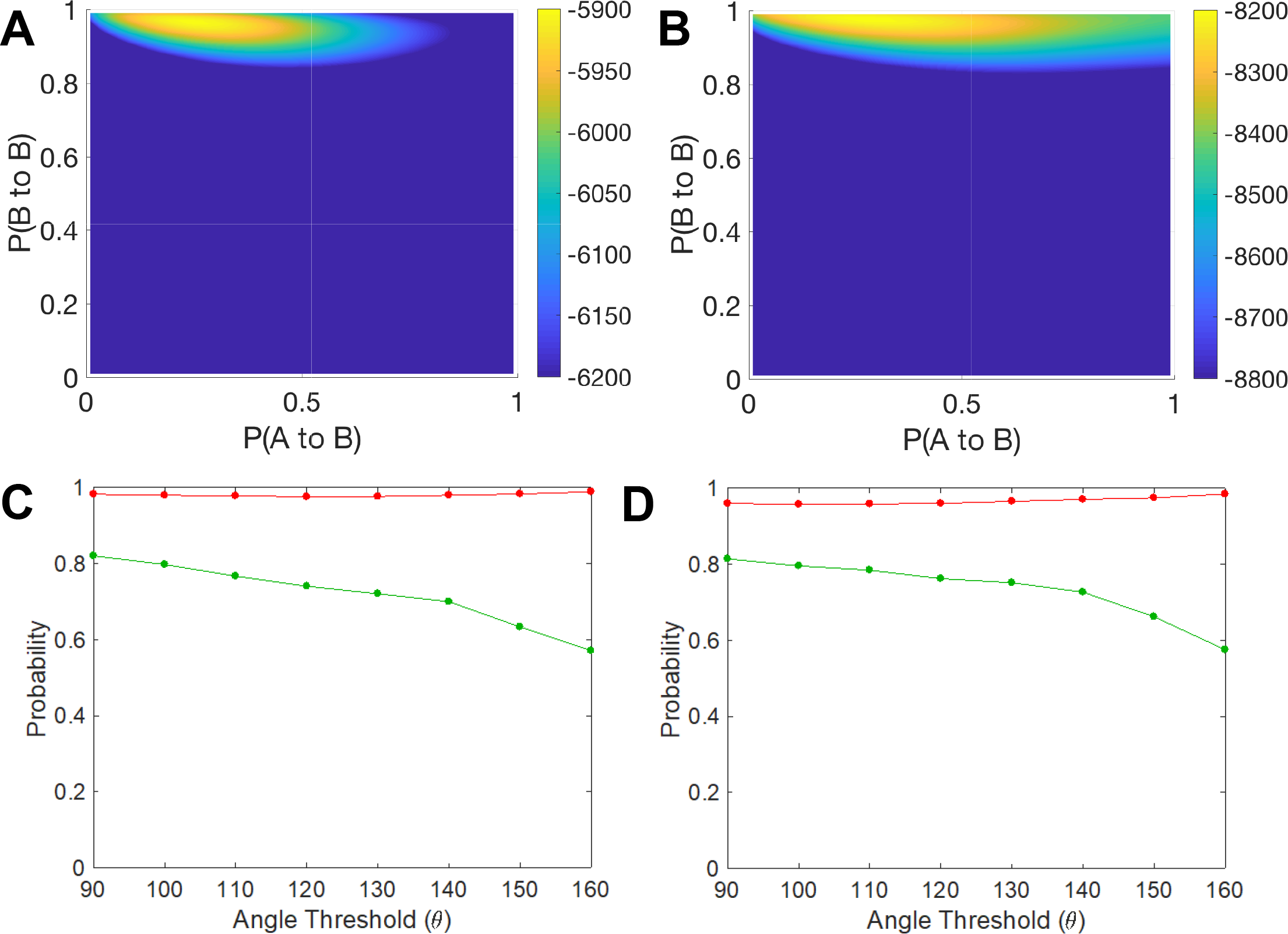
HMM sensitivity and convergence analysis. **A and B)** Log-likelihood of all the observed sequences of VAMP72-labeled vesicles (A) and myosin XIa (B) in protoplasts for an array of possible transition matrices. x and y-axes represent *P*(*h_t_* = *A* | *h*_*t*-1_ = *B*) and *P*(*h_t_* = *B* | *h*_*t*-1_ = *B*), respectively. Blue to yellow color map indicates log-likely hood where blue indicates less likely observed sequences and yellow indicates more likely sequences. A clear global maximum can be found in both (A) and (B). **C and D)** Hidden Markov Model myosin XIa (C) and VAMP72-labeled vesicle (D) transition probabilities in protoplasts, as a function of the angle threshold. Red lines indicate the Brownian to Brownian transition probability, *P*(*h*_*t*_ = *B* | *h*_*t*-1_ = *B*). Green lines indicate active to active transition probability, *P*(*h*_*t*_ = *A* | *h*_*t*-1_ = *A*).

## References

Avisar, D., Abu-Abied, M., Belausov, E. and Sadot, E. (2012). Myosin XIK is a major player in cytoplasm dynamics and is regulated by two amino acids in its tail. J Exp Bot 63, 241–9.

Avisar, D., Abu-Abied, M., Belausov, E., Sadot, E., Hawes, C. and Sparkes, I. A. (2009). A comparative study of the involvement of 17 *Arabidopsis* myosin family members on the motility of Golgi and other organelles. Plant Physiology 150, 700–9.

Avisar, D., Prokhnevsky, A. I., Makarova, K. S., Koonin, E. V. and Dolja, V. V. (2008). Myosin XI-K Is required for rapid trafficking of Golgi stacks, peroxisomes, and mitochondria in leaf cells of Nicotiana benthamiana. Plant Physiology 146, 1098–108.

Bibeau, J. P., Kingsley, J. L., Furt, F., Tuzel, E. and Vidali, L. (2018). F-Actin mediated focusing of vesicles at the cell tip is essential for polarized growth. Plant Physiol 176, 352–363.

Binder, B. and Holzhutter, H. G. (2012). A hypothetical model of cargo-selective rab recruitment during organelle maturation. Cell Biochem Biophys 63, 59–71.

Bove, J., Vaillancourt, B., Kroeger, J., Hepler, P. K., Wiseman, P. W. and Geitmann, A. (2008). Magnitude and direction of vesicle dynamics in growing pollen tubes using spatiotemporal image correlation spectroscopy and fluorescence recovery after photobleaching. Plant Physiology 147, 1646–1658.

Campàs, O. and Mahadevan, L. (2009). Shape and dynamics of tip-growing cells. Current Biology 19, 2102–2107.

Cole, R. A. and Fowler, J. E. (2006). Polarized growth: maintaining focus on the tip. Current Opinion in Plant Biology 9, 579–588.

de Graaf, B. H. J., Cheung, A. Y., Andreyeva, T., Levasseur, K., Kieliszewski, M. and Wu, H. M. (2005). Rab11 GTPase-regulated membrane trafficking is crucial for tip-focused pollen tube growth in tobacco. Plant Cell 17, 2564–2579.

Deschout, H., Martens, T., Vercauteren, D., Remaut, K., Demeester, J., De Smedt, S. C., Neyts, K. and Braeckmans, K. (2013). Correlation of dual colour single particle trajectories for improved detection and analysis of interactions in living cells. Int J Mol Sci 14, 16485–514.

Dumais, J., Shaw, S. L., Steele, C. R., Long, S. R. and Ray, P. M. (2006). An anisotropic-viscoplastic model of plant cell morphogenesis by tip growth. International Journal of Developmental Biology 50, 209–222.

Erickson, H. P. (2009). Size and Shape of Protein Molecules at the Nanometer Level Determined by Sedimentation, Gel Filtration, and Electron Microscopy. Biological Procedures Online 11, 32–51.

Fan, G. H., Lapierre, L. A., Goldenring, J. R., Sai, J. and Richmond, A. (2004). Rab11-family interacting protein 2 and myosin Vb are required for CXCR2 recycling and receptor-mediated chemotaxis. Molecular Biology of the Cell 15, 2456–69.

Fayant, P., Girlanda, O., Chebli, Y., Aubin, C. E., Villemure, I. and Geitmann, A. (2010). Finite element model of polar growth in pollen tubes. Plant Cell 22, 2579–93.

Furt, F., Lemoi, K., Tüzel, E. and Vidali, L. (2012). Quantitative analysis of organelle distribution and dynamics in *Physcomitrella patens* protonemal cells. BMC Plant Biology 12, 70.

Furt, F., Liu, Y. C., Bibeau, J. P., Tüzel, E. and Vidali, L. (2013). Apical myosin XI anticipates F-actin during polarized growth of *Physcomitrella patens* cells. Plant J 73, 417–28.

Geitmann, A. and Ortega, J. K. (2009). Mechanics and modeling of plant cell growth. Trends Plant Sci 14, 467–78.

Gilroy, S. and Jones, D. L. (2000). Through form to function: root hair development and nutrient uptake. Trends in Plant Science 5, 56–60.

Griffing, L. R., Gao, H. T. and Sparkes, I. (2014). ER network dynamics are differentially controlled by myosins XI-K, XI-C, XI-E, XI-I, XI-1, and XI-2. Front Plant Sci 5, 218.

Hammer, J. A. and Sellers, J. R. (2012). Walking to work: roles for class V myosins as cargo transporters. Nature Reviews Molecular Cell Biology 13, 13–26.

Heckman, D. S., Geiser, D. M., Eidell, B. R., Stauffer, R. L., Kardos, N. L. and Hedges, S. B. (2001). Molecular evidence for the early colonization of land by fungi and plants. Science 293, 1129–1133.

Henn, A. and Sadot, E. (2014). The unique enzymatic and mechanistic properties of plant myosins. Curr Opin Plant Biol 22, 65–70.

Hepler, P. K., Vidali, L. and Cheung, A. Y. (2001). Polarized cell growth in higher plants. Annual Review of Cell and Developmental Biology 17, 159–187.

Kang, M. and Kenworthy, A. K. (2008). A closed-form analytic expression for FRAP formula for the binding diffusion model. Biophysical Journal 95, L13–L15.

Kang, M. C., Day, C. A., DiBenedetto, E. and Kenworthy, A. K. (2010). A Quantitative Approach to Analyze Binding Diffusion Kinetics by Confocal FRAP. Biophysical Journal 99, 2737–2747.

Kenrick, P. and Crane, P. R. (1997). The origin and early evolution of plants on land. Nature 389, 33–39.

Kingsley, J. L., Bibeau, J. P., Mousavi, S. I., Unsal, C., Chen, Z., Huang, X., Vidali, L. and Tuzel, E. (2018). Characterization of cell boundary and confocal effects improves quantitative FRAP analysis. Biophys J 114, 1153–1164.

Konopka, C. A. and Bednarek, S. Y. (2008). Variable-angle epifluorescence microscopy: a new way to look at protein dynamics in the plant cell cortex. Plant Journal 53, 186–196.

Krementsov, D. N., Krementsova, E. B. and Trybus, K. M. (2004). Myosin V: regulation by calcium, calmodulin, and the tail domain. Journal of Cell Biology 164, 877–86.

Kroeger, J. H., Daher, F. B., Grant, M. and Geitmann, A. (2009). Microfilament orientation constrains vesicle flow and spatial distribution in growing pollen tubes. Biophys J 97, 1822–31.

Kroeger, J. H., Zerzour, R. and Geitmann, A. (2011). Regulator or Driving Force? The Role of Turgor Pressure in Oscillatory Plant Cell Growth. Plos One 6, -.

Lapierre, L. A., Kumar, R., Hales, C. M., Navarre, J., Bhartur, S. G., Burnette, J. O., Provance, D. W., Mercer, J. A., Bahler, M. and Goldenring, J. R. (2001). Myosin Vb is associated with plasma membrane recycling systems. Molecular Biology of the Cell 12, 1843–1857.

Li, J. F. and Nebenfuhr, A. (2008). The tail that wags the dog: the globular tail domain defines the function of myosin V/XI. Traffic 9, 290–8.

Li, X. D., Mabuchi, K., Ikebe, R. and Ikebe, M. (2004). Ca2+-induced activation of ATPase activity of myosin Va is accompanied with a large conformational change. Biochem Biophys Res Commun 315, 538–45.

Lise, M. F., Wong, T. P., Trinh, A., Hines, R. M., Liu, L. D., Kang, R. J., Hines, D. J., Lu, J., Goldenring, J. R., Wang, Y. T. et al. (2006). Involvement of myosin Vb in glutamate receptor trafficking. Journal of Biological Chemistry 281, 3669–3678.

Liu, Y. C. and Vidali, L. (2011). Efficient polyethylene glycol (PEG) mediated transformation of the moss *Physcomitrella patens*. J Vis Exp 50, e2560.

Loren, N., Hagman, J., Jonasson, J. K., Deschout, H., Bernin, D., Cella-Zanacchi, F., Diaspro, A., McNally, J. G., Ameloot, M., Smisdom, N. et al. (2015). Fluorescence recovery after photobleaching in material and life sciences: putting theory into practice. Quarterly Reviews of Biophysics 48, 323–387.

Luo, N., Yan, A., Liu, G., Guo, J., Rong, D., Kanaoka, M. M., Xiao, Z., Xu, G., Higashiyama, T., Cui, X. et al. (2017). Exocytosis-coordinated mechanisms for tip growth underlie pollen tube growth guidance. Nat Commun 8, 1687.

Madison, S. L., Buchanan, M. L., Glass, J. D., McClain, T. F., Park, E. and Nebenfuhr, A. (2015). Class XI myosins move specific organelles in pollen tubes and are required for normal fertility and pollen tube growth in Arabidopsis. Plant Physiology 169, 1946–1960.

Madison, S. L. and Nebenfuhr, A. (2013). Understanding myosin functions in plants: are we there yet? Curr Opin Plant Biol 16, 710–7.

McNally, J. G. (2008). Quantitative FRAP in analysis of molecular binding dynamics in vivo. Methods Cell Biol 85, 329–51.

Menand, B., Calder, G. and Dolan, L. (2007). Both chloronemal and caulonemal cells expand by tip growth in the moss *Physcomitrella patens*. J Exp Bot 58, 1843–9.

Nedvetsky, P. I., Stefan, E., Frische, S., Santamaria, K., Wiesner, B., Valenti, G., Hammer, J. A., Nielsen, S., Goldenring, J. R., Rosenthal, W. et al. (2007). A role of myosin Vb and Rab11-FIP2 in the aquaporin-2 shuttle. Traffic 8, 110–123.

Ohyama, A., Komiya, Y. and Igarashi, M. (2001). Globular tail of myosin-V is bound to VAMP/synaptobrevin. Biochemical and Biophysical Research Communications 280, 988–991.

Ojangu, E. L., Järve, K., Paves, H. and Truve, E. (2007). *Arabidopsis thaliana* myosin XIK is involved in root hair as well as trichome morphogenesis on stems and leaves. Protoplasma 230, 193–202.

Ovecka, M., Lang, I., Baluska, F., Ismail, A., Illes, P. and Lichtscheidl, I. K. (2005). Endocytosis and vesicle trafficking during tip growth of root hairs. Protoplasma 226, 39–54.

Park, E. and Nebenfuhr, A. (2013). Myosin XIK of Arabidopsis thaliana accumulates at the root hair tip and is required for fast root hair growth. Plos One 8, e76745.

Parton, R. M., Fischer-Parton, S., Watahiki, M. K. and Trewavas, A. J. (2001). Dynamics of the apical vesicle accumulation and the rate of growth are related in individual pollen tubes. J Cell Sci 114, 2685–95.

Peremyslov, V. V., Klocko, A. L., Fowler, J. E. and Dolja, V. V. (2012). *Arabidopsis* myosin XI-K localizes to the motile endomembrane vesicles associated with F-actin. Frontiers in Plant Science 3, 184.

Peremyslov, V. V., Prokhnevsky, A. I., Avisar, D. and Dolja, V. V. (2008). Two class XI myosins function in organelle trafficking and root hair development in *Arabidopsis*. Plant Physiology 146, 1109–1116.

Peremyslov, V. V., Prokhnevsky, A. I. and Dolja, V. V. (2010). Class XI myosins are required for development, cell expansion, and F-Actin organization in *Arabidopsis*. Plant Cell 22, 1883–97.

Preuss, M. L., Serna, J., Falbel, T. G., Bednarek, S. Y. and Nielsen, E. (2004). The *Arabidopsis* Rab GTPase RabA4b localizes to the tips of growing root hair cells. Plant Cell 16, 1589–1603.

Prokhnevsky, A. I., Peremyslov, V. V. and Dolja, V. V. (2008). Overlapping functions of the four class XI myosins in *Arabidopsis* growth, root hair elongation, and organelle motility. Proceedings of the National Academy of Sciences, USA 105, 19744–19749.

Pruyne, D. W., Schott, D. H. and Bretscher, A. (1998). Tropomyosin-containing actin cables direct the Myo2p-dependent polarized delivery of secretory vesicles in budding yeast. Journal of Cell Biology 143, 1931–45.

Rabiner, L. R. (1989). A tutorial on Hidden Markov-Models and selected applications in speech cecognition. Proceedings of the Ieee 77, 257–286.

Roding, M., Guo, M., Weitz, D. A., Rudemo, M. and Sarkka, A. (2014). Identifying directional persistence in intracellular particle motion using Hidden Markov Models. Mathematical Biosciences 248, 140–145.

Rodriguez, O. C. and Cheney, R. E. (2002). Human myosin-Vc is a novel class V myosin expressed in epithelial cells. Journal of Cell Science 115, 991–1004.

Rojas, E. R., Hotton, S. and Dumais, J. (2011). Chemically mediated mechanical expansion of the pollen tube cell wall. Biophys J 101, 1844–53.

Roland, J. T., Kenworthy, A. K., Peranen, J., Caplan, S. and Goldenring, J. R. (2007). Myosin Vb interacts with Rab8a on a tubular network containing EHD1 and EHD3. Molecular Biology of the Cell 18, 2828–2837.

Rybak, K., Steiner, A., Synek, L., Klaeger, S., Kulich, I., Facher, E., Wanner, G., Kuster, B., Zarsky, V., Persson, S. et al. (2014). Plant cytokinesis is orchestrated by the sequential action of the TRAPPII and exocyst tethering complexes. Developmental Cell 29, 607–20.

Schott, D., Ho, J., Pruyne, D. and Bretscher, A. (1999). The COOH-terminal domain of Myo2p, a yeast myosin V, has a direct role in secretory vesicle targeting. Journal of Cell Biology 147, 791–807.

Sparkes, I. A., Teanby, N. A. and Hawes, C. (2008). Truncated myosin XI tail fusions inhibit peroxisome, Golgi, and mitochondrial movement in tobacco leaf epidermal cells: a genetic tool for the next generation. Journal of Experimental Botany 59, 2499–2512.

Sprague, B. L. and McNally, J. G. (2005). FRAP analysis of binding: proper and fitting. Trends Cell Biol 15, 84–91.

Szumlanski, A. L. and Nielsen, E. (2009). The Rab GTPase RabA4d regulates pollen tube tip growth in *Arabidopsis thaliana*. Plant Cell 21, 526–544.

Szymanski, D. B. and Cosgrove, D. J. (2009). Dynamic coordination of cytoskeletal and cell wall systems during plant cell morphogenesis. Curr Biol 19, R800–11.

Tominaga, M., Kimura, A., Yokota, E., Haraguchi, T., Shimmen, T., Yamamoto, K., Nakano, A. and Ito, K. (2013). Cytoplasmic streaming velocity as a plant size determinant. Developmental Cell 27, 345–52.

Ueda, H., Yokota, E., Kutsuna, N., Shimada, T., Tamura, K., Shimmen, T., Hasezawa, S., Dolja, V. V. and Hara-Nishimura, I. (2010). Myosin-dependent endoplasmic reticulum motility and F-actin organization in plant cells. Proc Natl Acad Sci U S A 107, 6894–9.

Uemura, T., Ueda, T., Ohniwa, R. L., Nakano, A., Takeyasu, K. and Sato, M.H. (2004). Systematic analysis of SNARE molecules in Arabidopsis: dissection of the post-Golgi network in plant cells. Cell Struct Funct 29, 49–65.

van Gisbergen, P. A., Li, M., Wu, S. Z. and Bezanilla, M. (2012). Class II formin targeting to the cell cortex by binding PI(3,5)P(2) is essential for polarized growth. Journal of Cell Biology 198, 235–50.

Vick, J. K. and Nebenfuhr, A. (2012). Putting on the breaks: regulating organelle movements in plant cells(f). J Integr Plant Biol 54, 868–74.

Vidali, L., Augustine, R. C., Kleinman, K. P. and Bezanilla, M. (2007). Profilin is essential for tip growth in the moss *Physcomitrella patens.* Plant Cell 19, 3705–22.

Vidali, L., Burkart, G. M., Augustine, R. C., Kerdavid, E., Tüzel, E. and Bezanilla, M. (2010). Myosin XI is essential for tip growth in *Physcomitrella patens*. Plant Cell 22, 1868–82.

Vidali, L., Rounds, C. M., Hepler, P. K. and Bezanilla, M. (2009). Lifeact-mEGFP reveals a dynamic apical F-actin network in tip growing plant cells. Plos One 4, e5744.

Volpicelli, L. A., Lah, J. J., Fang, G., Goldenring, J. R. and Levey, A. I. (2002). Rab11a and myosin Vb regulate recycling of the M4 muscarinic acetylcholine receptor. J Neurosci 22, 9776–84.

Wang, F., Thirumurugan, K., Stafford, W. F., Hammer, J. A., 3rd, Knight, P.J. and Sellers, J. R. (2004). Regulated conformation of myosin V. J Biol Chem 279, 2333–6.

Yan, Q., Sun, W., Kujala, P., Lotfi, Y., Vida, T. A. and Bean, A. J. (2005). CART: An hrs/actinin-4/BERP/myosin V protein complex required for efficient receptor recycling. Molecular Biology of the Cell 16, 2470–2482.

Zonia, L. and Munnik, T. (2008). Vesicle trafficking dynamics and visualization of zones of exocytosis and endocytosis in tobacco pollen tubes. Journal of Experimental Botany 59, 861–873.

